# The unbiased estimation of the fraction of variance explained by a model

**DOI:** 10.1101/2020.10.30.361253

**Authors:** Dean A. Pospisil, Wyeth Bair

## Abstract

The correlation coefficient squared, *r*^2^, is often used to validate quantitative models on neural data. Yet it is biased by trial-to-trial variability: as trial-to-trial variability increases, measured correlation to a model’s predictions decreases; therefore, models that perfectly explain neural tuning can appear to perform poorly. Many solutions to this problem have been proposed, but some prior methods overestimate model fit, the utility of even the best performing methods is limited by the lack of confidence intervals and asymptotic analysis, and no consensus has been reached on which is the least biased estimator. We provide a new estimator, 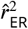, that outperforms all prior estimators in our testing, and we provide confidence intervals and asymptotic guarantees. We apply our estimator to a variety of neural data to validate its utility. We find that neural noise is often so great that confidence intervals of the estimator cover the entire possible range of values ([0, 1]), preventing meaningful evaluation of the quality of a model’s predictions. We demonstrate the use of the signal-to-noise ratio (SNR) as a quality metric for making quantitative comparisons across neural recordings. Analyzing a variety of neural data sets, we find ~ 40% or less of some neural recordings do not pass even a liberal SNR criterion.

**Author Summary:** Quantifying the similarity between a model and noisy data is fundamental to the verification of advances in scientific understanding of biological phenomena, and it is particularly relevant to modeling neuronal responses. A ubiquitous metric of similarity is the correlation coefficient. Here we point out how the correlation coefficient depends on a variety of factors that are irrelevant to the similarity between a model and data. While neuroscientists have recognized this problem and proposed corrected methods, no consensus has been reached as to which are effective. Prior methods have wide variation in their precision, and even the most successful methods lack confidence intervals, leaving uncertainty about the reliability of any particular estimate. We address these issues by developing a new estimator along with an associated confidence interval that outperforms all prior methods.

## Introduction

Building an understanding of the nervous system requires the quantification of model performance on neural data. It is typical to estimate Pearson’s *r*^2^ of neural responses with a model fit as such a quantification. Yet the typical estimator, 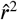, is fundamentally confounded by the trial-to-trial variability of neural responses: a low *r*^2^ could be the result of a poor model or high neuronal variability.

One approach to this problem is to average over many repeated trials of the same stimulus in order to reduce the influence of trial-to-trial variability. With a finite number of trials this approach will never wholly remove the influence of noise and its confounding effect, moreover, the collection of additional trials is expensive. A more principled approach has been to account for trial-to-trial variability in the estimation of the fraction of explainable variance or *r*^2^. Most often this takes the form of attempting to estimate what the *r*^2^ would have been in the absence of trial-to-trial variability. Here we call this quantity 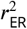, the *r*^2^ between the model prediction and the expected response (ER) of the neuron (i.e., the ‘true’ mean, or expected value, of the tuning curve). While a variety of solutions have been proposed to estimate this quantity (Roddey, Girish, and Miller, 2000; Pasupathy and Connor, 2001; Sahani and Linden, 2003; Hsu et al., 2004; David and Gallant, 2005; Haefner and Cumming, 2009; Yamins et al., 2012; Schoppe et al., 2016), they have not been quantitatively compared thus there is no basis to reach a consensus on which methods are appropriate, or more importantly inappropriate. We find that several estimators still in recent use have large biases. Moreover, estimators that had relatively small biases did not have associated confidence intervals, thus the degree of uncertainty in these sometimes highly variable estimates remains ambiguous. Finally, none of these methods have been analyzed asymptotically to give a theoretical guarantee that they will converge to 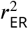, in other words it has not been shown that they are consistent estimators.

To address these substantial problems, we introduce 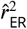, which is a simple analytic estimator of 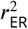, along with a method for generating *α*-level confidence intervals. We validate our estimator in simulation, prove that it is consistent, and provide head-to-head comparisons to prior methods. We then demonstrate the use of 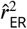 and its confidence interval on two sets of neural data. We find many cases where neuronal data is so noisy that estimates of 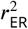 provide little inferential power about the quality of a model fit. This naturally leads to a useful metric of the quality of a neuronal recording that we will refer to as the signal-to-noise ratio (SNR), and which can be interpreted in terms of the number of trials needed to reliably detect tuning. Across a diverse set of neural recordings, we find that many neurons do not pass even a liberal criterion for providing meaningful insight into the quality of a model fit.

## Results

The results of the paper are organized as follows. We first give the essential intuition into the source of the bias in 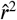 and how 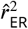 removes this bias. Next, we evaluate 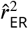 through simulation and compare it to prior methods. We then demonstrate the method on two neural data sets: one from a study of motion direction tuning in area MT and one from a study of responses to natural images in area V4. Finally, we develop the signal-to-noise ratio (SNR) as a metric to determine the inferential power of a given neuronal recording.

Consider a typical scenario in sensory neuroscience where the responses of a neuron to *m* stimuli across *n* repeated trials of each stimulus have been collected and the average of these responses, the estimated tuning curve (Figure 1, dashed green line), is compared to responses predicted by a model (red line). Even if the *m* expected values of the neuronal response, *μ_i_*, (solid green trace), perfectly correlate with the model predictions, *v_i_* (red trace is scaled and shifted relative to green), the *m* sample averages, *Y_i_* (dashed green trace), will deviate from their expected value owing to the sample mean’s variability. Here, we quantify this variability as the variance, *σ*^2^, of the distribution of responses from trial-to-trial (see Methods). We assume *σ*^2^ is constant across responses to different stimuli given a variance stabilizing transform. The variance of the sample mean across all stimuli will thus be 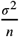. Owing to the variance of the sample mean, the reported 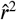 can be appreciably less than 1 even though the *r*^2^ between the underlying expected values of the neuronal response and the model is 1.

**Figure 1.**
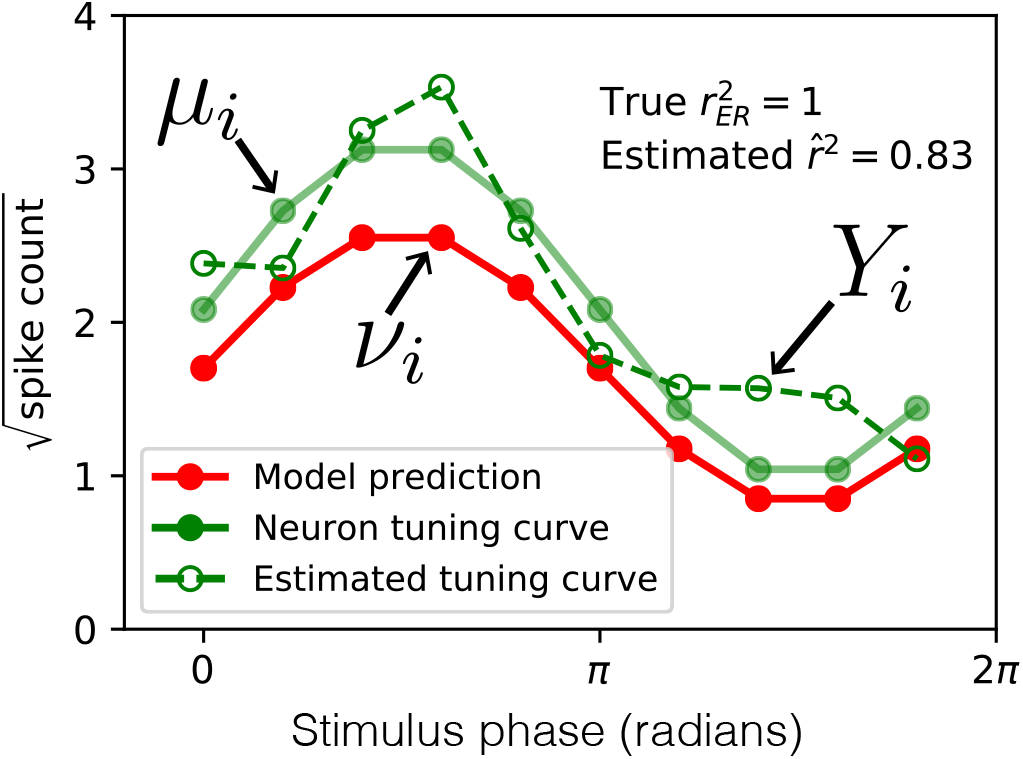
Sampling noise confounds estimation of the correlation between model prediction and neuronal tuning curve. The expected (true) spike count in response to a set of 10 stimuli (solid green trace) is perfectly correlated with a model (red trace), yet owing to sampling error (neural trial-to-trial variability) the estimated tuning curve (green dashed curve) has correlation less than one with the model 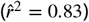.

The quantity we attempt to estimate in this paper is *r*^2^ between the model predictions (*v_i_*) and the expected neuronal responses (*μ_i_*). We will call this quantity 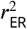, the fraction of variance of the ‘expected response’ explained:

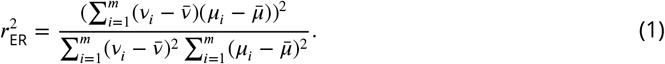

We will show that the naive sample estimator, which uses *Y_i_* in place of *μ_i_*,

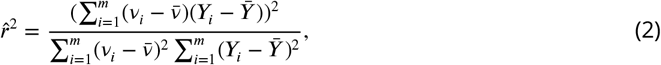

has an expected value that can be well approximated as the ratio of the expected values of its numerator and denominator as follows (for asymptotic justification see Methods, “Inconsistency of 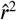 in *m*”):

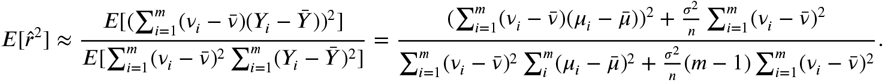

While the terms on the left in the numerator and denominator are the same as 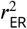, the terms on the right are proportional to the trial-to-trial variability (*σ*^2^) and cause 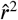 to deviate from 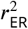. This is the essential problem: 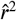 is biased away from 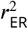 proportional to the amount of variability, 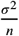, in the estimated responses.

The strategy we take to solve this problem is straightforward: find unbiased estimators of these noise terms and subtract them from the numerator and denominator of Eqn. 2 for 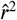:

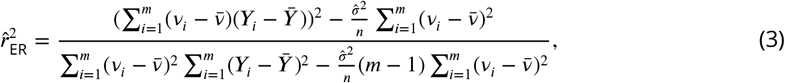

where 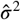 is an unbiased estimator for trial-to-trial variability, after a variance stabilizing transform if necessary. Typically 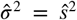, the sample variance, but not necessarily. For example if stimuli are shown only once (*n* = 1), trial-to-trial variability could be assumed and substituted into 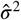. The numerator and denominator of the fraction 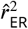 are unbiased estimators of the numerator and denominator of 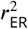. This solution is approximate since the expected value of a ratio is not necessarily the ratio of the expected values of the numerator and denominator. Yet we show in simulation the approximation is very good for typical neural statistics, and we show analytically that, unlike 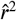, our estimator 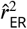 converges to the true 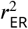 as the number of stimuli *m* → ∞ (see Methods, “Consistency of 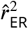 in *m*”). We next evaluate this estimator in simulation.

### Validation of 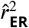 by simulation

To demonstrate the effectiveness and key properties of 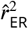, we ran a simulation with *m* = 362 stimuli, *n* = 4 repeats, and *σ*^2^ = 0.25 (the trial-to-trial variance of Poisson neuronal response after a variance-stabilizing transform, see Methods: Assumptions and terminology for derivation of unbiased estimator). This corresponds, for example, to presenting 362 different shape stimuli for 4 repeats each, which is seen as a minimal amount of data to characterize shape tuning in V4 neurons, yet would take on the order of 1 hour.

In the case where the model prediction (*v_i_*) and expected response (*μ_i_*) were perfectly correlated (as in Figure 1) and SNR was moderate at 0.5 (i.e., the neuron was substantially modulated by the stimuli), the distribution of the naive estimator, 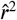, is centered well below 1 (Figure 2A, blue distribution). Thus, the model appears to be a poor fit to data that it in fact generated, indicating that 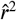 is a poor estimator of 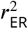. On the other hand, our corrected estimator, 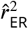, is appropriately centered at 1 (orange distribution). Approximately 50% of the time our estimator exceeds 1, taking on impossible values of 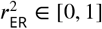, but this is necessary to achieve unbiased estimates for high 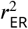 because truncating the values would shift the mean below 1.

**Figure 2.**
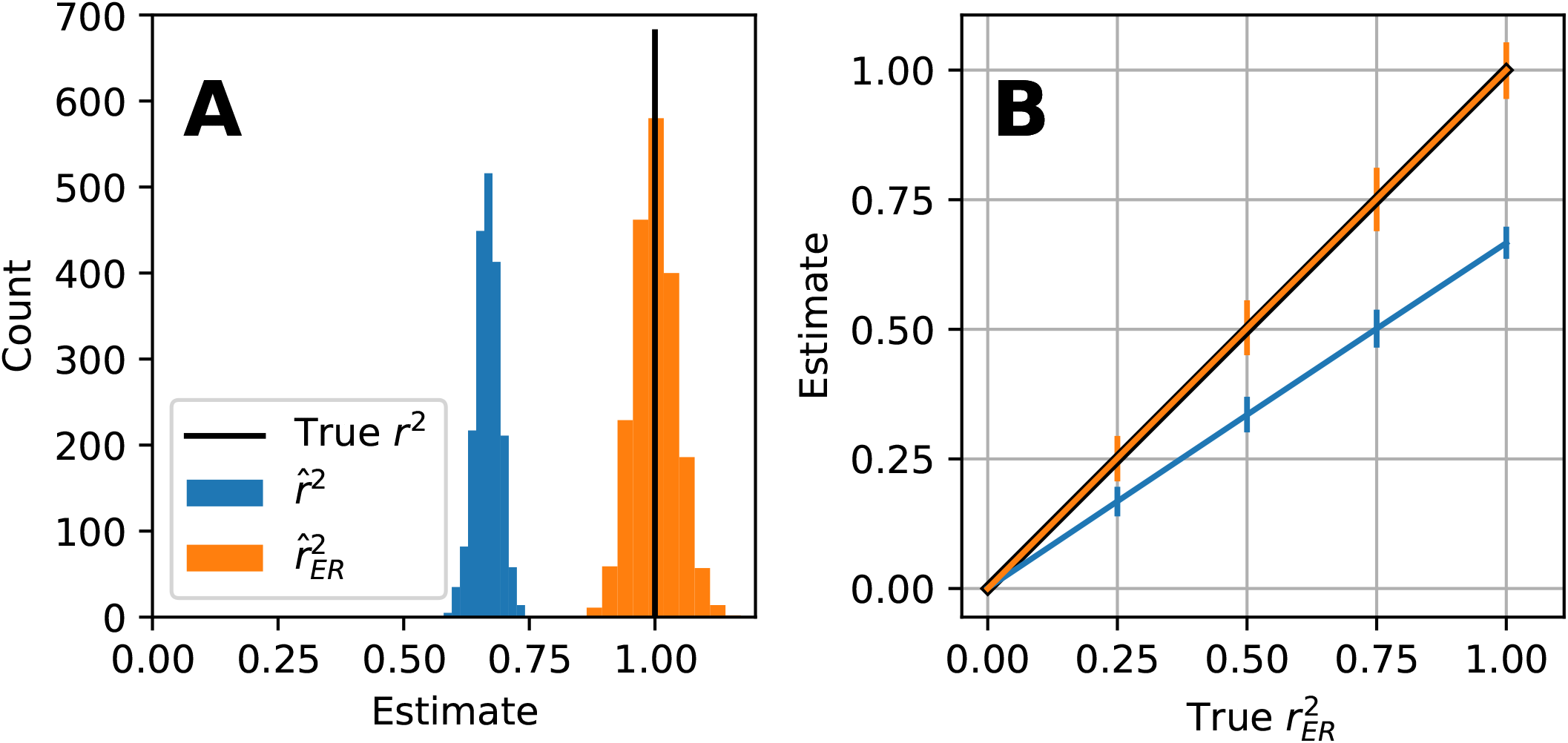
Simulation of 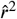 (blue) and 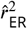 (orange) for estimating fit of model to neuron at varying levels of 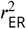 where m=362, *n* = 4, and *σ*^2^ = 0.25. **(A)** Simulation at lower dynamic range SNR = 0.5, even when *r*^2^ = 1, 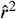 (blue) is on average 0.67 whereas 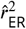 (orange) is on average 1.0002. The bias of 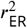 (see Methods: Bias of 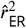) is small relative to its variability (95% quantile = [0.93, 1.07] vertical bars) and the bias of 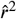. **(B)** Repeating the same simulation across levels of 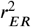 [0, 0.25, 0.5, 0.75, 1] there is appreciable bias present between true 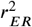 (black trace), and naive 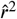 (blue) but not 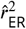 (orange trace).

We evaluated the estimators 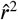 and 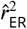 at five values of 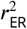 (0, 0.25, 0.5, 0.75, 1) and plotted the mean and 95% quantiles. Figure 2B shows that 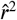 (blue line) grossly underestimates 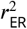 (black line) at all levels except for 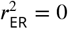, whereas 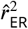 (orange line) correctly estimates the true correlation 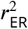 (orange and black line overlap). Thus the estimator 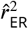 performs favorably in this simulation. Next, we characterize the performance of 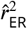 relative to 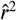 in simulations that cover a wide range of the parameters, *m*, *n* and SNR.

### Asymptotic properties of 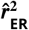 and 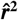

We ran simulations to determine the bias and variance of 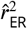 relative to 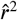 as a function of the parameters SNR, *n*, and *m*. Figure 3A shows that as SNR increases, 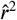 (blue) and 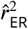 (orange) converge (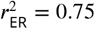, *n* = 4, *m* = 362). Thus, for neuronal recordings with strong stimulus modulation, these two estimators should have similar values. At low values of SNR, e.g., 0.1, 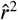 has a large downward bias (mean 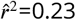), whereas 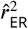 has a small upward bias relative to its own variability and to the bias of 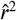 (for the source of this bias see Methods: Bias of 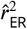). This small upward bias of 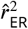 quickly diminishes as SNR increases, whereas the large negative bias of 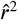 remains across a much wider range of SNR. The essential problem this simulation reveals is that if SNR varies widely from neuron to neuron, the bias in the naive estimate will cause apparent variation in *r*^2^ across neurons that depends on SNR and not on the underlying neural function. Neuronal SNR is not typically under experimental control, making this problem difficult to avoid.

**Figure 3.**
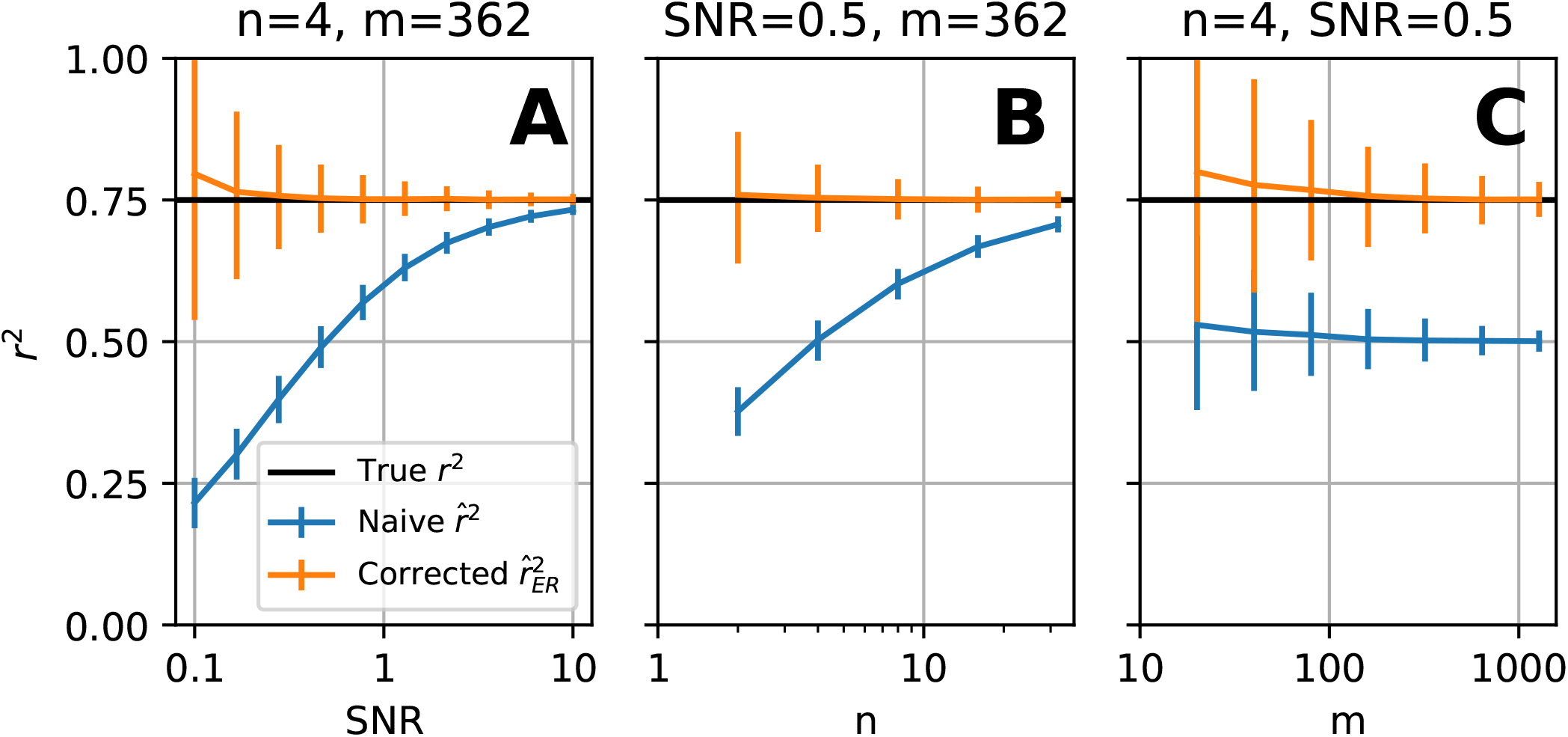
Simulation of 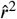 (blue) and 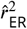 (orange) for estimating fit of model to neuron at varying levels of SNR, *n*, and *m*. **(A)** Relationship of naive 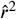 and corrected 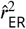 with SNR. Simulating when 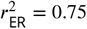 (horizontal black line), *m* = 362, *n* = 4, and *σ*^2^ = 0.25 we evaluated the estimators 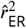 and 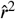 (respectively orange and blue) average and 95% confidence intervals. **(B)** Relationship of estimators with *n* the number of repeats. **(C)** Relationship of naive 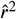 and corrected 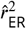 with *m* the number of stimuli. In this simulation all parameters but *m* which is varied and SNR=0.5 which is fixed are the same as A.

The number of repeats, *n*, is under the experimenter’s control but is expensive to increase. Figure 3B shows how 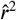 and 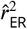 converge as *n* increases. Thus the bias in 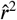 can be reduced by increasing the number of repeats, but to achieve this requires a very high number of repeats for low SNR. An advantage of 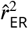 is that even for low *n* it on average estimates the true correlation to the model (orange trace overlaps black trace, Figure 3B) providing a large gain in total trial efficiency for estimating the quality of model fit.

When increasing the number of stimuli, *m*, unlike the previous two cases, 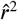 and 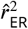 do *not* converge to the same value (Figure 3C). While variability of both estimators decreases (error bars become smaller), it is clear in simulation that 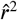 is not a consistent estimator of 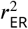 in *m* since it does not converge to 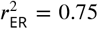. While there is a small upward bias of 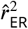 for low *m*, as *m* increases this bias is reduced (see Methods, “Consistency of 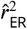 in *m*”).

### Comparison to prior methods

Accounting for noise when interpreting the fit of models to neural data has been examined and applied in the neuroscientific literature for some time (Roddey, Girish, and Miller, 2000; Pasupathy and Connor, 2001; Sahani and Linden, 2003; Hsu et al., 2004; David and Gallant, 2005; Haefner and Cumming, 2009; Yamins et al., 2012; Schoppe et al., 2016). Several studies have followed the approach of attempting to estimate the upper bound on the quality of fit of a model given noise and then referencing the measure of fit to this quantity. Roddey et al. estimate this upper bound by computing their estimate of model fit, ‘coherence’, across split trials then plotting coherence of the data to the model predictions relative to the split trial coherence. Yamins et al. normalize *r*^2^ with split-trial correlation transformed by the Spearman-Brown prediction formula (averaged across randomly resampled subsets of trials); we will call this 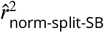. Hsu et al. also use split-half correlation (averaged across randomly resampled subsets of trials), to estimate an upper bound (CC_max_) by a transformation they derive attempting to estimate the correlation of the true mean with the firing rate of the neuron. For purpose of comparison, we square this estimator and call it 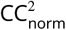. Kindel et al. measure the correlation of a Gaussian simulation based on the sample mean and variance of the neural data with the sample mean as a measure of CC_max_. They then normalize the Pearson’s *r* between model predictions and neural responses with CC_max_; we call this estimator 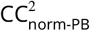 (PB for parametric bootstrap). Pasupathy and Connor estimate the fraction of total variance accounted for by trial-to-trial variability, intuitively the fraction of unexplainable variance, then use it to normalize *r*^2^. We call this estimator 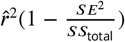. With a similar motivation Cadena et al (2019) provide a metric they call “fraction explainable variance explained” (FEVE). They form the ratio of mean squared prediction error over total variance of the response except subtracting off an estimate of trial-to-trial variability from both. They then subtract this ratio from one. While all of these methods might be intuitively appealing, the quantities to which they converge, and their relationship to 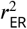 is unclear. Here, instead, we follow a line of research (Sahani and Linden, 2003; Haefner and Cumming, 2009) that explicitly attempts to construct an unbiased estimator of *r*^2^ in the absence of noise (see Discussion: Relationship to prior methods). Specifically, they calculate an approximate expectation of their estimator and then adjust the estimator to remove its bias. Sahani and Linden derive an an estimator they call ‘normalized signal power explained’ but do not specify the form of the model being fit under which the estimator behaves reasonably (intercept and slope linear regression, see Appendix: Normalized Signal Power Explained (SPE)). While Haefner and Cumming’s (2008) 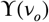 takes inspiration from Sahani and Linden, they specify that the model being fit is linear in its coefficients and appropriately take into account degrees of freedom. Heretofore many of the methods reviewed above have not been quantitatively validated and none have been directly compared, thus there is little basis upon which to select one over the other.

We now compare all these methods with respect to their MSE in estimating 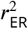 (see Methods: Simulation procedure). We exclude from this comparison David and Gallant (2005) because their method depends on a large number of repeated trials, at which point the estimators’ utility decreases. Choosing *n* = 4 trials and *m* = 362 stimuli, we sort the estimators (Figure 4 y-axis) by their MSE in a test case where 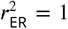. We generally find, 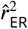, SPE_norm_, 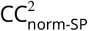, FEVE and 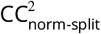 are all comparable in their performance (red trace, top 6 points) with 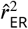 performing slightly, but significantly, better. SPE_norm_ and 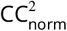 are numerically identical in their performance and their trial-to-trial results. On the other hand 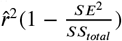 and 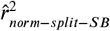 both grossly over estimate 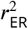 and the naive estimator 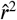 as expected under estimates (mean=0.50). In addition 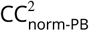 grossly underestimates 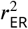. When the true 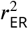 is 0.5 we find similar results where 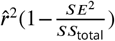 and 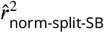 respectively on average overestimate (0.63 and 1.04 respectively) and mean 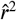 is 0.25. Thus, serious caution should be taken when interpreting these last two estimators. We conclude 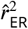 is as good as any estimator of 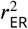 available, has a simple analytic form, and can still be calculated without calculating the sample variance, for example if no repeats are collected and variance must be assumed. None of the top six estimators we reviewed have associated confidence intervals and thus we now provide confidence intervals for 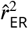.

**Figure 4.**
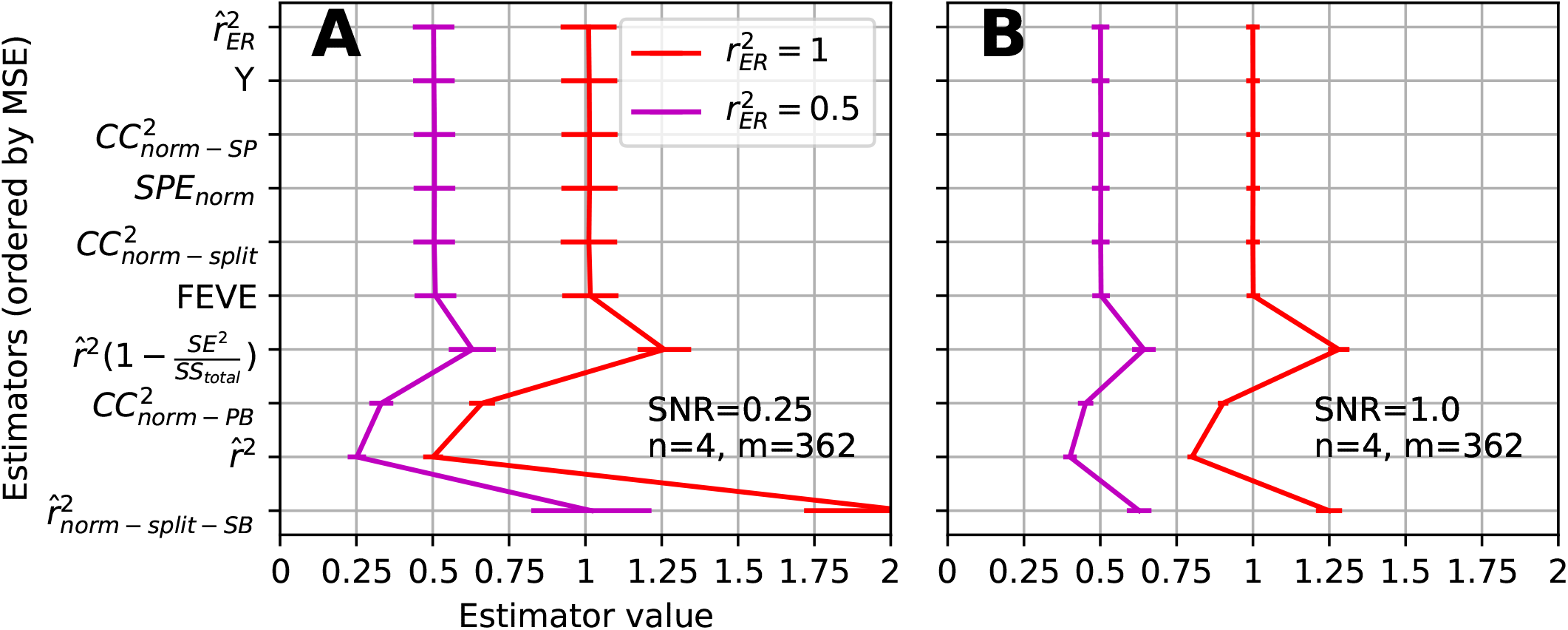
Comparison of estimators of 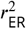 on the basis of simulated data (see Fig 2A). **(A)** Low SNR simulation where estimators on vertical axis are sorted from top to bottom by smallest MSE with respect to estimating 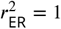 (red trace shows mean and SD of each estimator). **(B)** Same simulation at higher SNR (1.0) but same *m*, *n*.

### Confidence intervals for 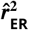

In order to interpret point estimates such as 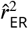, it is important to be able to meaningfully quantify uncertainty about the estimate relative to the true parameter 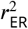. An *α*-level confidence interval (CI) provides an interval that will contain the true parameter *α* × 100% of the time for IID estimates. We considered three typical generic approaches to forming CIs for 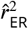: the non-parametric bootstrap, the parametric bootstrap, and BC_*α*_ (Efron and Tibshirani, 1994). We found all methods to be lacking because they did not achieve the desired *α* in simulations with ground truth. Motivated by these problems, we developed a Bayesian method we call the ‘Estimate-Centered Credible Interval’ (ECCI). We first recount the issues we found with the bootstrap methods and then provide a basic account of the Bayesian method we use throughout the paper. For more detailed exposition, see Methods: Quantifying uncertainty in the estimator.

The non-parametric bootstrap is a commonly used method to approximate CIs. In our case, it involves randomly re-sampling with replacement from the *n* trials in response to each of the *m* stimuli then calculating 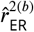 for the bootstrap sample. Repeating this many times allows the quantiles of the bootstrap distribution of 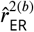 to be used as CIs. We applied this method across a simulated population of 3000 neurons with *m* = 40 and *n* = 4 and found it suffered from two problems. First, the CIs were not centered around 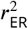, specifically the interval was too low (Figure 5A), with the upper and lower bounds of the interval (orange and blue traces, respectively) almost always falling below the true value (green). Secondly, as the true 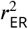 increased from 0 to 1, CIs contained 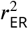 at rates far lower than the desired *α* = 0.8 (Figure 6 cyan trace, open-circles under the trace indicate a significant difference, *p* < 0.01 Bonferoni corrected z-test). Thus at practically all levels of correlation, the non-parametric bootstrap performs poorly. The problem is a result of the expected value of the empirical distribution (the sample mean) being typically much lower than 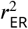. To overcome this, we turned to the parametric bootstrap where we could explicitly estimate 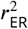 with our unbiased estimator. This method approximates CIs by estimating the parameters of an assumed distribution from which samples are generated. In our case it involves estimating *σ*^2^, *d*^2^ and 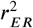 and then simulating observations from the distribution with these parameters to calculate 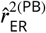. Drawing many 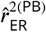 we again use the sample quantiles as CI estimates. Figure 5B shows that this overcomes the main failure of the non-parametric bootstrap, but this method tended to be too conservative for low 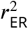 values (Figure 6 red trace below 0.8 at left side) and too liberal for high values (red trace above 0.8 at right side). Deviations such as these are well known for bootstrap percentile methods when the variance is a non-constant function of the mean and/or the distribution of the estimator is skewed (Efron and Tibshirani, 1994). The correction to the bootstrap, the BC_*α*_ (bias-corrected and accelerated) bootstrap, can help ameliorate these issues by implicitly approximating the skewness and the mean-variance relationship from bootstrap samples. We employed BC_*α*_ with our parametric bootstrap and found that performance improved but still deviated from the desired *α* for low and high 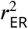 (Figure 6, green trace).

**Figure 5.**
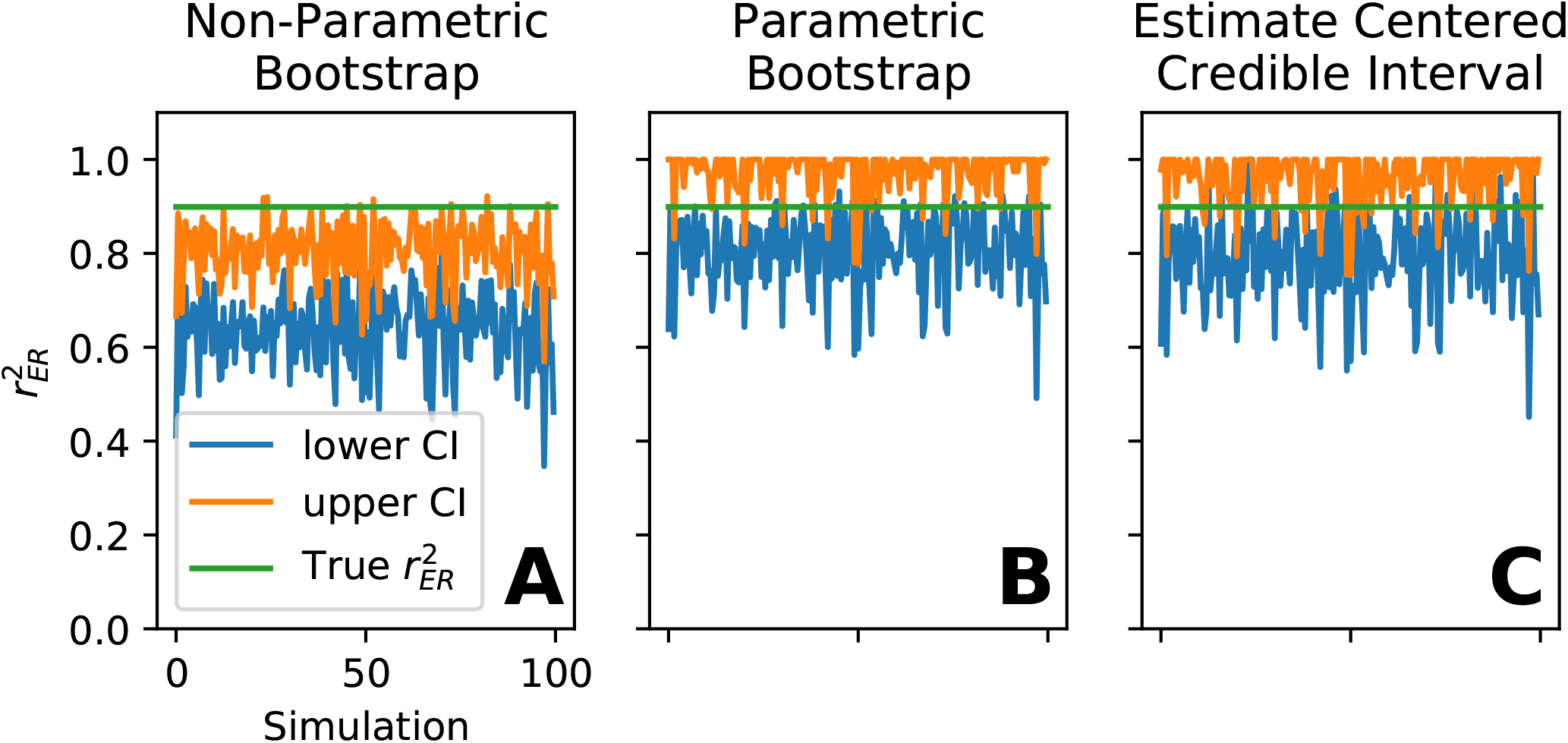
Example plots of confidence interval methods simulations. Parameters of simulation were *n* = 4, *m* = 40, and 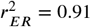, dynamic range (*d*^2^ = 0.25) and trial-to-trial variability (*σ*^2^ = 0.25) and the target confidence level was *α* = 0.8. There were 2000 simulations, 200 of which are plotted here. Confidence intervals for each method were calculated for the same set of responses. **(A)** Simulation of non-parametric bootstrap confidence interval, true correlation value indicated by horizontal green line. The upper end of the confidence interval (orange) and lower end (blue) are not centered around the true correlation and the green line is not contained within the confidence interval far more often than 1 - *α* = 0.2. For quantification see Figure 6 blue trace. **(B)** Simulation of non-parametric bootstrap confidence interval. **(C)** Simulation of estimate centered credible interval method.

**Figure 6.**
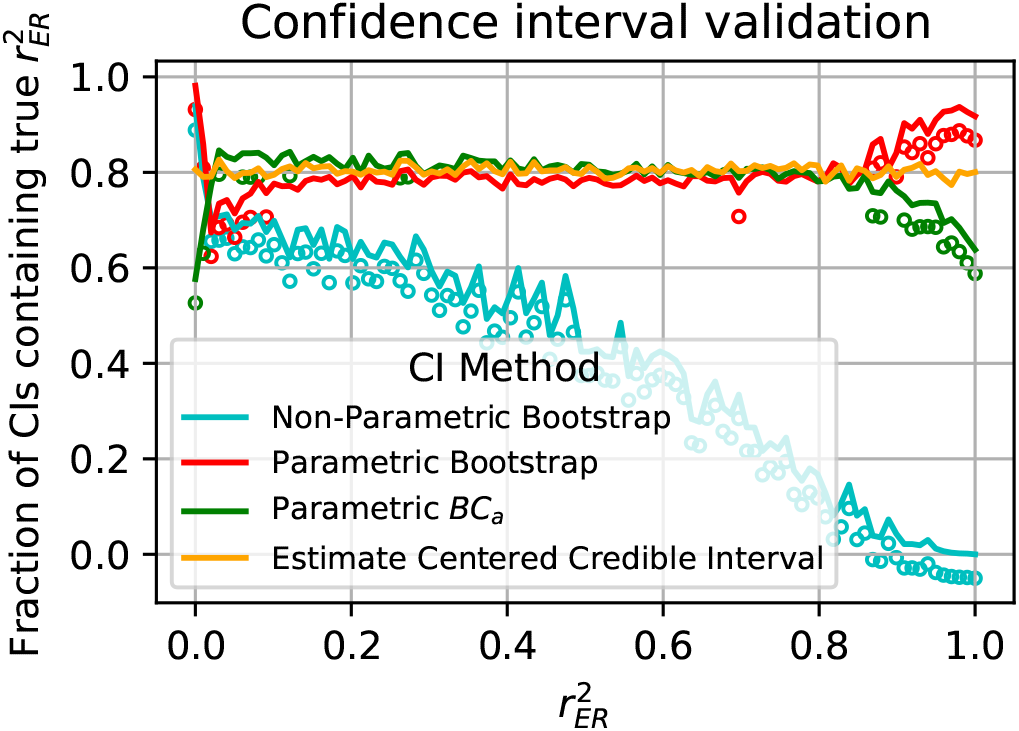
Results of confidence interval simulation. The fraction of times the confidence interval contained the true value are plotted for each method (see Legend) as a function of the true correlation value 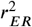 at 100 values linearly spaced between 0 and 1. If the fraction intervals deviated from 0.8 significantly (p<0.01 Bonferroni corrected across all tests (400)) it is indicated with an open circle.

We aimed to create a CI with better *α*-level performance. To do this, we assumed uninformative priors on the parameters *σ*^2^ and *d*^2^ so that, conditioned on estimates of these parameters, we can draw from the distribution of 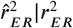 for an arbitrary 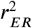 (see Methods: Confidence Intervals for 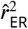). This allows us to compute the highest true 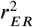 that would have given an observed 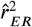 or a lowervalue in *α*/2 proportion of IID samples. We take this as the high end, 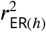, of our CI. Similarly we determine the low end, 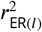 of the CI as the lowest 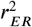 that produces a value greater than or equal to 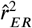 in *α*/2 of the samples. In Methods we give conditions under which this procedure will provide *α* level CIs (see Methods: Confidence intervals for 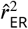). In our simulations this method consistently achieves the desired *α* at all levels of 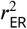 (Figure 6, orange trace). We use this CI method, which we call the estimate-centered credible interval (ECCI), throughout the rest of the paper.

### Application of estimator to MT data

We have shown in simulation that the use of 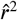 introduces ambiguity as to whether a low correlation value was the result of trial-to-trial variability or a poor model, whereas 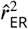 removes this ambiguity. Here we demonstrate in neural data how this, in tandem with confidence intervals, allows investigators to distinguish between units that systematically deviate from model predictions versus those that simply have noisy responses. We re-analyzed data from single neurons in area MT responding to dot motion in eight equally spaced directions (Zohary et al., 1994; Bair et al., 2001). A classic model of these responses is a single cycle sinusoid as a function of the direction of dot motion with the free parameters phase, amplitude, and average firing rate. We chose this MT experiment as the first example application because the data has a high number of repeats (*n* = 10) and a low dimensional model, thus it is simple to visually inspect whether the neuronal tuning curves agree with the model predictions.

An example of a typical MT neuron direction tuning curve (Figure 7A, orange trace) has an excellent sinusioidal fit (blue trace), as reflected in its estimated 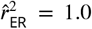. Furthermore, the short confidence interval ([0.99, 1.0]) quantifies the lack of ambiguity about the quality of the fit. On the other hand, the tuning curve of a neuron with 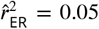 (Figure 7B) has a clear systematic deviation from the least-squares fit. Here the tuning curve is double peaked and thus largely orthogonal to any single cycle sinusoid. It is important to notice that this neuron has far lower SNR (SNR=3.2), thus without 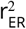 there would be plausible doubts about whether the correlation was lower because of SNR or systematic deviation. Furthermore, with low SNR it would be plausible that the estimate itself is noisy (Figure 3A), but the short confidence interval ([0.01,0.1]) characterizes the fit as being systematically poor.

**Figure 7.**
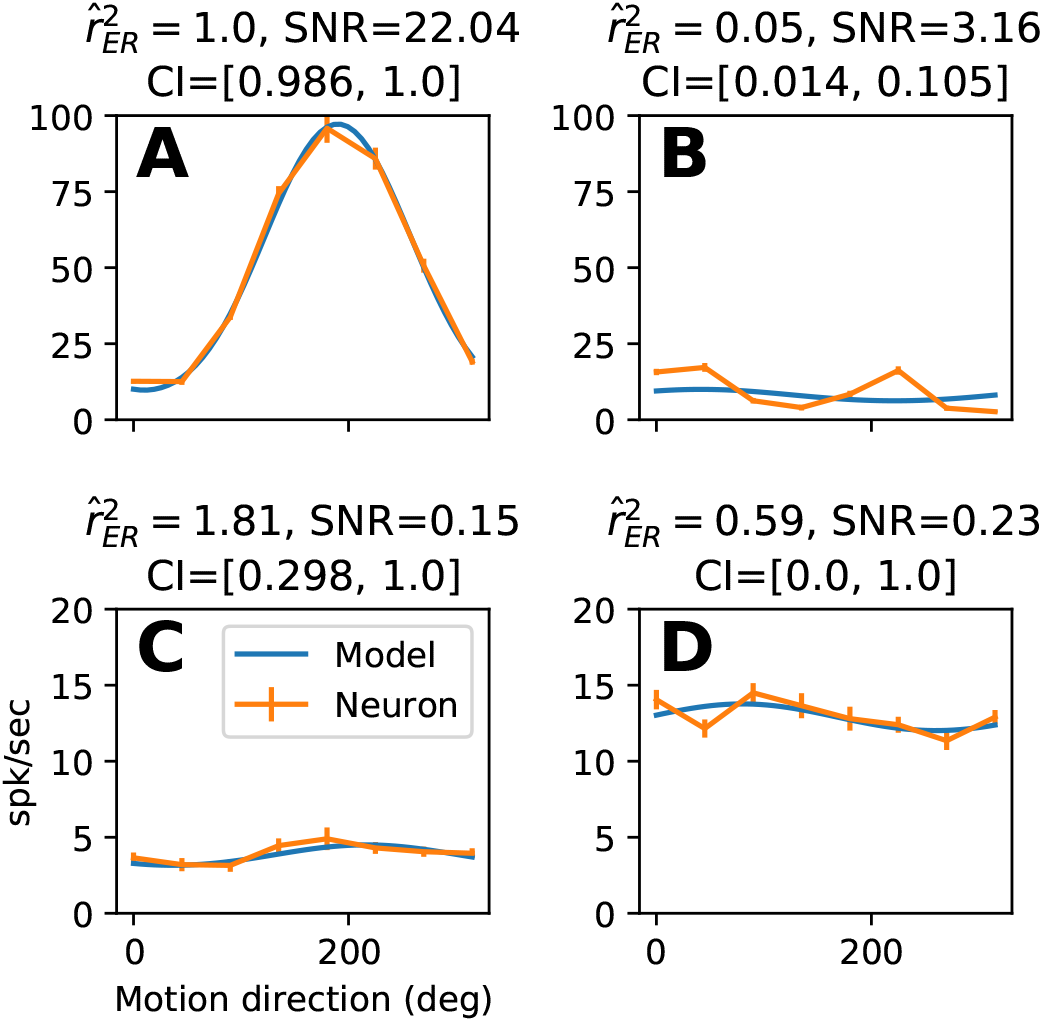
Example figures of sinusoidal model fit to MT data. **(A)** Example cell (orange trace with SEM bars) with near perfect fit to sinusoidal model (blue trace). **(B)** Example cell with poor fit to sinusoidal model but with reasonable SNR. **(C)** Example cell with poor SNR and wild estimate of 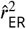. Poor SNR is reflected in CI covering entire interval [0,1] **(D)** Example cell with reasonable seeming 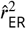 estimate but low SNR reflected in CI covering entire interval [0,1] thus estimate cannot be trusted.

In some cases, neurons show little tuning for direction and thus have very low SNR over a set of directional stimuli. This in turn can cause 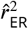 to give wild estimates (Figure 7C, SNR=0.15, 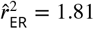). If we truncate the value to the nearest possible 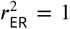, we might be tempted to interpret this as a well-fit direction selective neuron. But, the CI covers most of the interval of possible values ([0.3, 1]), making it clear that little information can be gleaned about 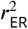 from this data. Extreme 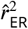 values themselves can indicate when the estimator is unreliable, but even a reasonable seeming 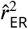 value, for example 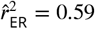 (Figure 7D), can be unreliable when there is a low SNR (0.23). In this case, the confidence interval covers the maximal range ([0,1]), indicating that the point estimate is unreliable. Thus, 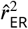 and its associated confidence interval quickly and unambiguously show how well the model fits the MT data, avoiding the tiresome and unreliable process of judging each fit by eye for the 162 neurons.

While we have shown to a good approximation that 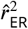 is unbiased and its expected value is largely invariant to SNR, this is definitely not the case for the variance of the estimator. In Figure 3A it is clear that the variability of the estimator is larger for lower SNR. This fact should be kept in mind when interpreting the spread of 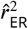 values. For example, we calculated 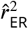 and confidence intervals across our entire population of MT neurons. Of the estimates with high SNR (Figure 8, right side, SNR > 4), most neurons are well fit to the model and only a few have less than 3/4 of their variance accounted for (8/81). For the estimates with lower SNR (SNR<4), left side of Figure 8), this fraction is substantially higher (39/81), but we must be wary that the increased variability of these estimates will spread out the distribution. When estimating population dispersion, conclusions may be confounded by the SNR-dependence of the variability of 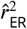.

**Figure 8.**
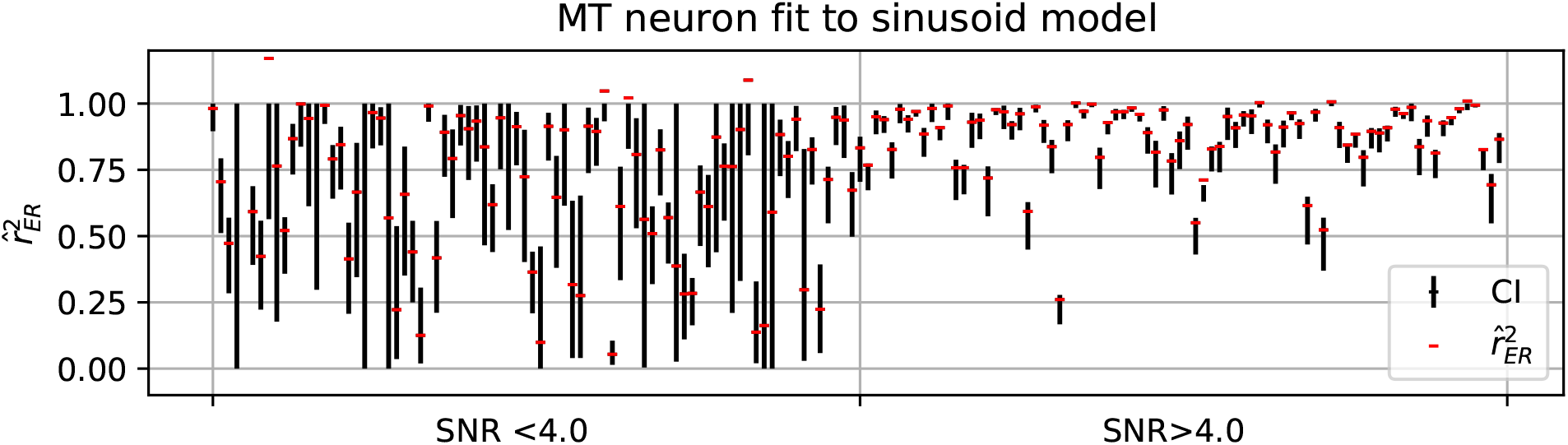
Confidence intervals (vertical black lines *α* = 90%) and point estimates (horizontal red lines) 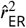 across all MT neurons fit to sinusoidal model. Estimates in first interval (left) had SNR less than the median (4), whereas in the second interval greater than the median.

Comparing 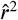 and 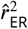 for the high SNR units, we find they have very similar estimates (Figure 9 most red points lie on diagonal). Thus one could exchange the two estimates and come to similar general conclusions about model fits. Yet the utility of 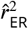 is not restricted to when it diverges from 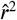, its utility is that it removes ambiguity about whether trial-to-trial variability may be spuriously pushing fits down. The interpretation of the naive estimator 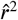 remains ambiguous until it can be confirmed it does not suffer from this issue.

**Figure 9.**
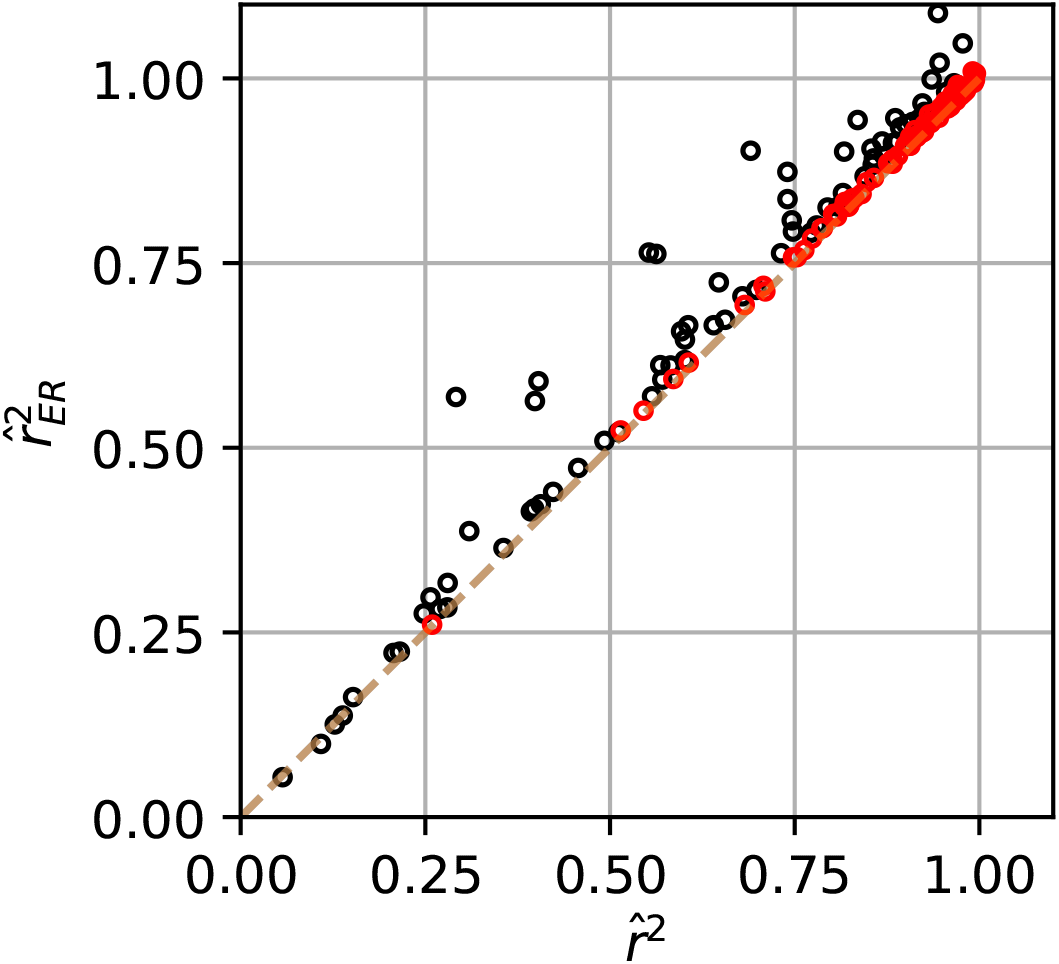
Relationship of naive 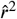 and corrected 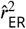 between fits of sinusoidal model to MT data. Units with SNR greater than the median across the population (SNR=4) are plotted in red and those below in black.

The MT data has only a few stimuli and many repeats but other experiments use a larger number of stimuli and, consequently, a lower number of repeats. Below we apply 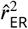 in these more challenging conditions.

### Application of estimator to V4 data

The primate mid-level visual cortical area V4 is known to have complex high-dimensional selectivity. To rigorously assess models of neuronal responses in areas like V4, validation needs to be performed on responses to a large corpus of natural images to ensure that models capture ecologically valid selectivity (Rust and Movshon, 2005). Thus, the number of unique stimuli, *m*, will be large at the expense of having relatively few repeats, *n*, and SNR can be low because stimuli are not customized to the preferences of a given neuron. These are the challenging conditions under which 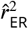 avoids the major confounds of 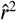. Here we estimate 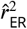 and associated 90% confidence intervals for a model that won the University of Washington V4 Neural Data Challenge (see Methods: Electrophysiological data) by most accurately predicting single-unit (SUA) and multi-unit activity (MUA) for held-out stimuli. Plotting 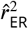 against 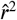 (Figure 10) shows that the corrected estimates are slightly higher than the naive estimates (points lie above diagonal line). Using 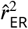 here is important because it provides confidence that the poor fit quality is not a result of noise and that the best performing model often did not explain more than 50% of the variance in the tuning curve.

**Figure 10.**
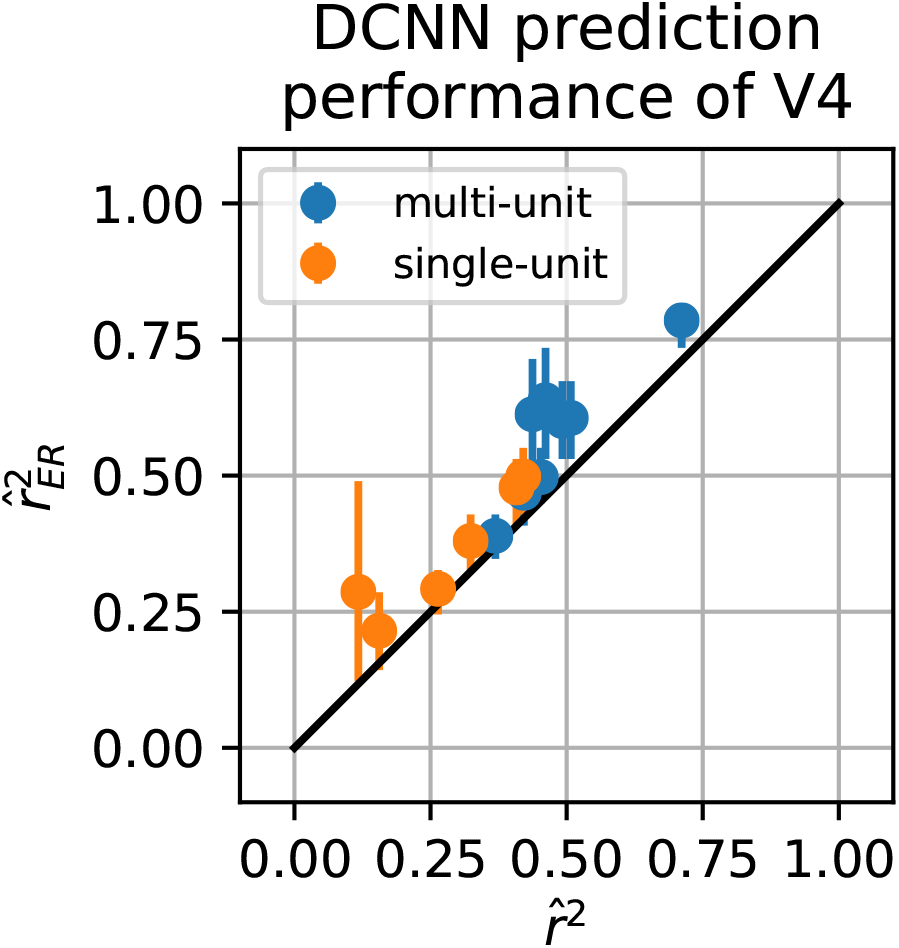
Performance of deep convolution neural network (DCNN) in predicting mean response of V4 to natural images. 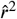 on x-axis vs 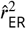 on y-axis show similar estimates though 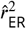 consistently reports slightly higher performance.

While we have examined 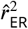 for individual recordings, it can also be useful to estimate the average quality of model fit across a population of neurons. Since the individual estimates are unbiased, the group average is also an unbiased estimate of the population mean 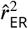. We computed such group means for the single-unit and multi-unit V4 recordings (Figure 11), and found that the model performed significantly better in predicting the responses of multi-unit activity (Welch’s t-test p=0.005, MUA mean=0.57, single unit mean=0.35). If instead the naive 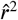 were used, this finding could have been dismissed as the result of multi-unit data having higher SNR and thus naturally higher 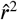. As it stands, this interesting observation can be followed up to potentially gain insight about the structure of selectivity across multiple units recorded nearby in V4.

**Figure 11.**
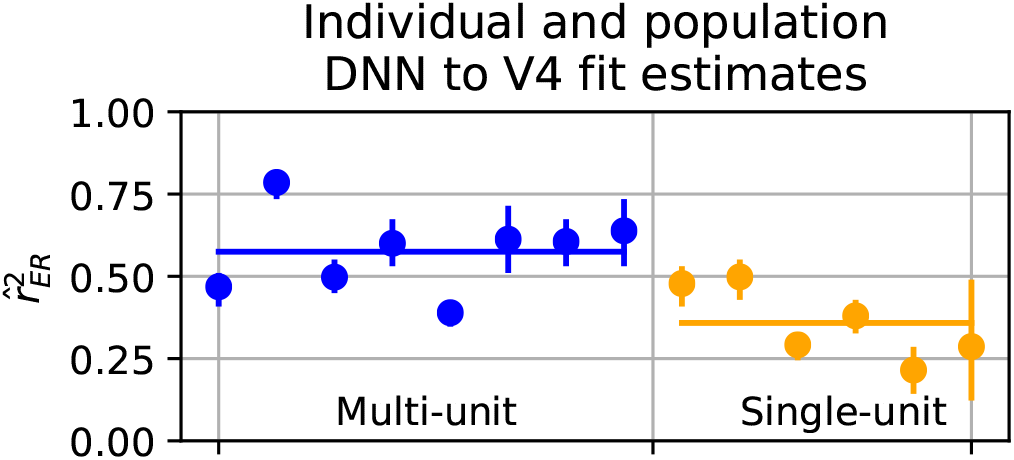
Comparison of single (orange) vs multi-unit (blue) fits with respect to individual estimated model fits and average. Multi-unit responses on average were better fit by the DNN then single units (Welch’s t-test t=3.7, p=0.005). Since individual estimates asymptotically unbiased the group average inherits this unbiasedness.

Finally, this V4 data set provides a good example of how using 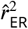 can allow testing a larger stimulus space, as predicted by simulations above in Figure 3B. Figure 12 shows that with 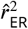 (solid lines) on average even two trials is enough to estimate the true correlation while the naive estimator requires more repeats (higher *n*) to converge. For example, for recording 1 (red), 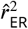 (solid line) on average predicts the same quality of model fit for two or more stimulus repeats, whereas even after six repeats the naive 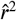 has not converged.

**Figure 12.**
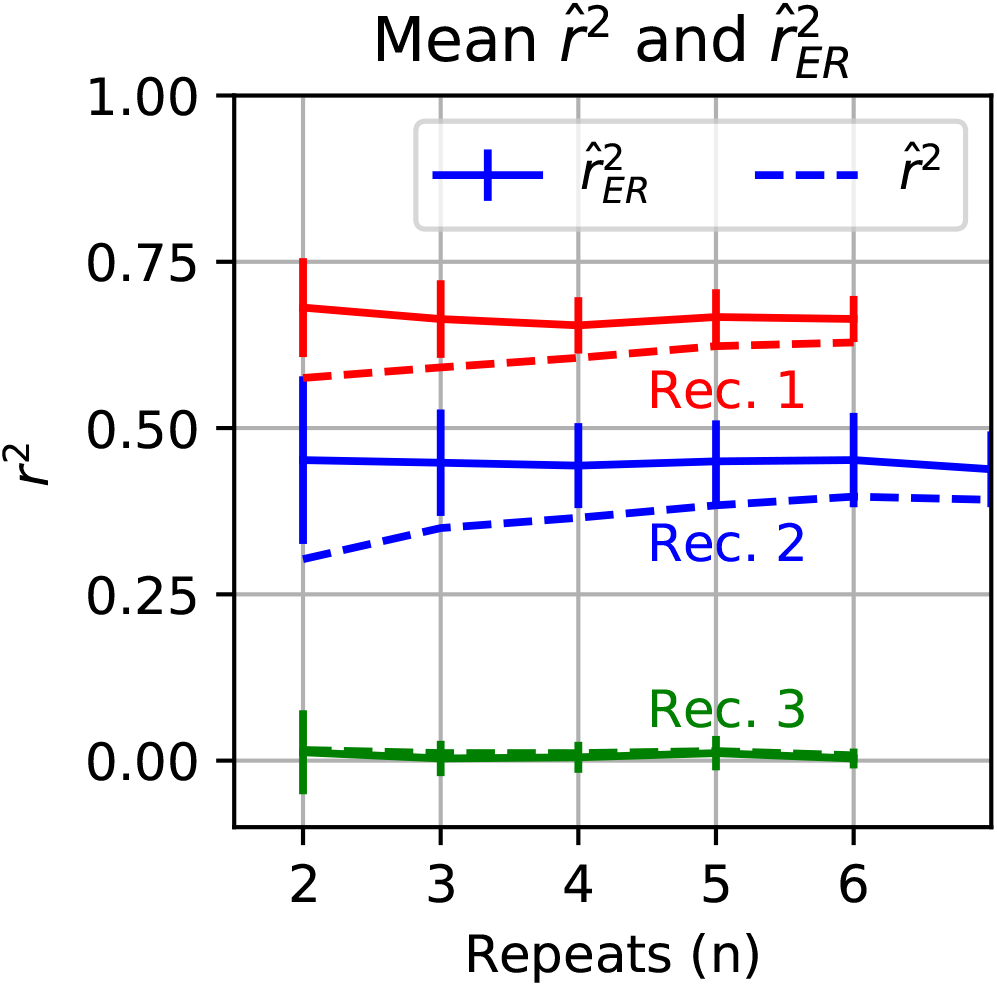
Relationship of naive 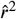 and corrected 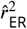 with *n* the number of trials for V4 data. Different colors indicate different recordings and solid lines are the average 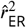 estimate across random with replacement shuffling of trials with vertical bars indicating the standard deviation and naive 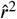 averages are the dashed lines.

### Signal-to-noise ratio as recording quality metric

We define the SNR for a neuronal tuning curve to be the ratio of the variation in the expected response across stimuli to the trial-to-trial variability across repeats:

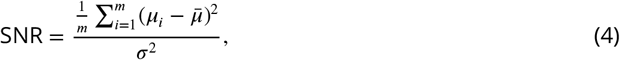

where *μ_i_* is the expected response to the *i*th stimulus and 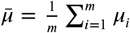. If SNR is close enough to 0, then no reasonable number of unique stimuli (*m*) or repeats (*n*) will allow successful inference about any non-flat tuning of the neuron. SNR is not a function of *n* or *m*, but it can vary across neurons, sets of stimuli and recording modalities. The expected range of SNR, based on prior data, should be taken into account when choosing *n* and *m* to achieve a criterion level of statistical power for an experiment. If SNR is high, one may save recording time by keeping *n* and *m* low. For experimental data, we do not know *μ_i_* in Eqn. 4, and rather than substituting sample estimates, *Y_i_*, which would give an inflated estimate, we use an equation that corrects for trial-to-trial noise (Eqn. 15, Methods).

We saw that the variability, and thus confidence interval length, of 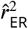 depended strongly on SNR (Figure 3A). In some neural data with low SNR, the CI covered the entire possible range of 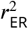 values, [0,1] (Figure 8, left side). This suggests that SNR can be employed as a simple metric to judge data quality across different organisms, recording modalities and brain regions, for the purpose of making comparative analyses and setting aside data that has little or no power. We first perform SNR comparisons across data sets, and then provide an interpretation of SNR in terms of the numbers of trials needed to reliably detect selectivity for the stimulus.

We examined a diverse collection of neural data sets (see Methods: Electrophysiological data) and found wide variation in SNR both within and across the data sets (Figure 13A). At the low end, calcium imaging data from cortical neurons in area VISp of mouse responding to gratings (pink trace *N* = 40,520 neuronal ROIs, Allen Brain Observatory, 2016) had a median SNR of 0.01, while at the high end, MT neurons in response to dot motion (Zohary et al., 1994) had a median SNR of 4.0 (blue trace, *N* = 162). A stimulus protocol nearly identical to that used for the VISp Ca^2+^ data (pink and gray traces for gratings and natural images, respectively) was used to collect the Allen Institute NeuroPixel electrode data (purple and brown traces *N* = 2,015); however, the Ca^2+^ data had a substantially lower SNR (0.01 and 0.02) compared to the electrode data (SNR 0.12 and 0.19), suggesting that this difference relates to the recording modality.

**Figure 13.**
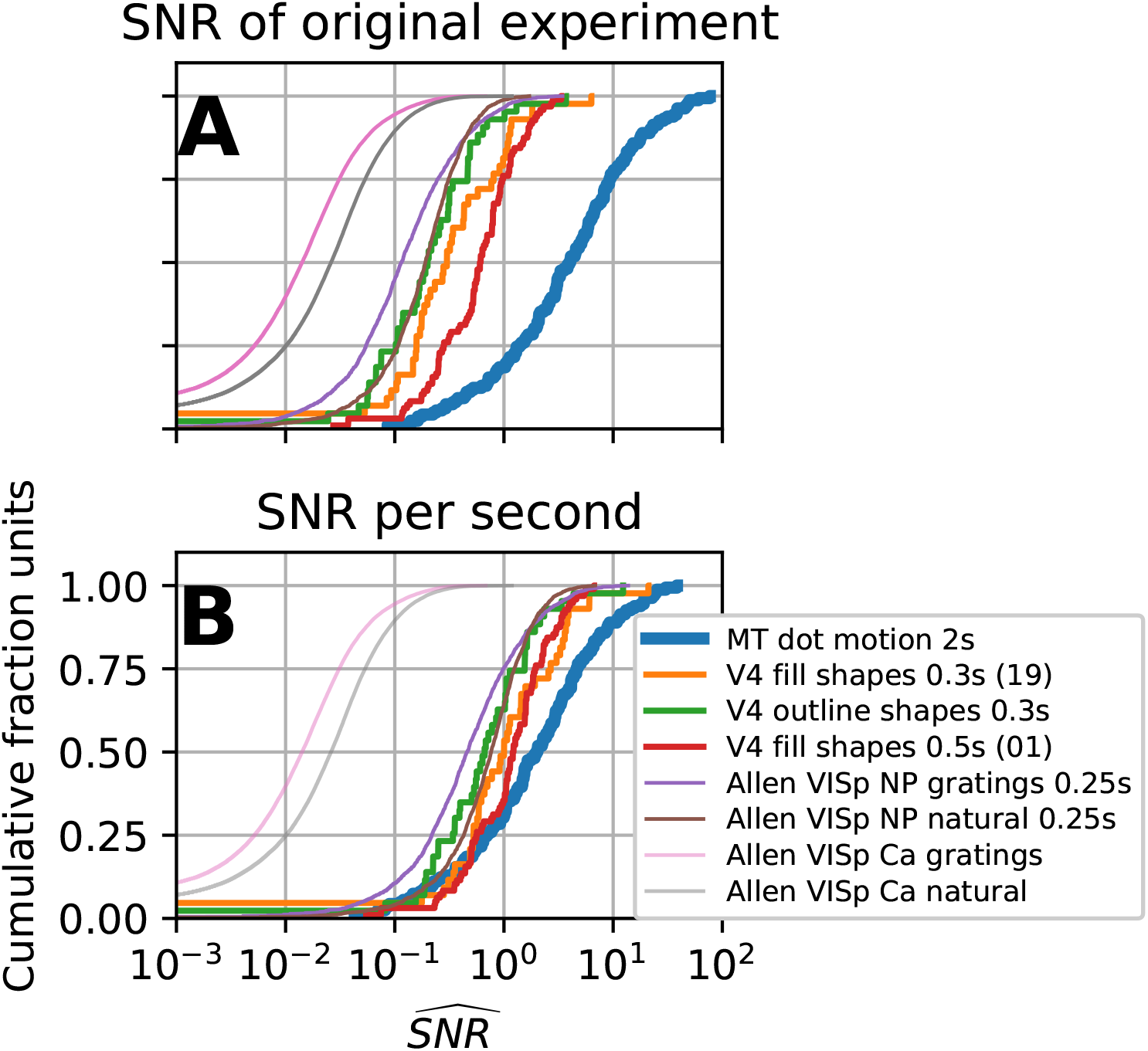
The cumulative distribution of signal-to-noise ratio (SNR) for several datasets. **(A)** The cumulative distribution of SNR given the original experiments protocol. Traces with the same line thickness have similar numbers of *n* and *m* (MT thick *n* ≈ 10, *m* = 8, V4 medium *n* ≈ 5, *m* ≈ 350, and Allen thin *n* ≈ 50 *m* ≈ 120). The Allen data has two recording modalities spiking recordings on Neuropixel probes (NP) and two-photon calcium imaging (Ca) both recorded for the same stimuli: natural scenes and gratings (see Methods: Electrophysiological data). **(B)** Distribution of SNR normalized with respect to spike counting window (except for calcium signal). Assuming original average spike count is same as the average spike rate across a 1 second window these traces represent the distribution of SNR for a 1 second spike counting window.

In the case of spiking neurons, SNR can be improved by increasing the stimulus duration and thus the spike counting window. Under the generally optimistic assumption that spike rate stays constant in the counting window, we can normalize SNR across the data-sets to what the SNR would have been had all spike count windows been 1 second long (Figure 13B). This reduces the differences in SNR across the spiking data-sets (the six right-most traces), thus the outstanding SNR of the MT data-set could potentially have been achieved if spike count windows had been longer for the other experiments. Still, of the spiking data, the Allen Neuropixel data has the lowest medians, thus additional efforts to ameliorate low SNR (via number of trials or stimulus choice) could be utilized. Furthermore, the assumption of a constant spike rate will hold to different degrees: neural responses can peak shortly after stimulus onset and then return close to baseline. Thus, different experimental conditions call for different standards for number of trials and stimulus duration to adequately characterize a tuning curve.

To provide concrete meaning to SNR, we suggest interpreting it in terms of the number of trials (*m* and *n*) needed to reliably detect stimulus modulation in an *F*-test. Specifically, for a given *m* and *n* we computed the minimal SNR required to achieve a high probability (*β* = 0.99) of rejecting the null hypothesis that the mean response to all stimuli is the same (see Methods: SNR relation to *F*-test and number of trials, Eqn. 21). We plot a color map of this minimal SNR as a function of *m* and *n* (Figure 14), where the diagonal grey contour lines indicate fixed total number of trials (*mn*) for different *m* : *n* ratios. In general, as the total number of trials increases (moving perpendicular to the grey diagonals toward the upper right), the SNR required for reliable tuning curve estimates decreases. The SNR threshold is lower when *n* is favored over *m* for the same number of total trials, i.e., the SNR threshold level iso-contours have steeper slopes than the grey diagonals.

**Figure 14.**
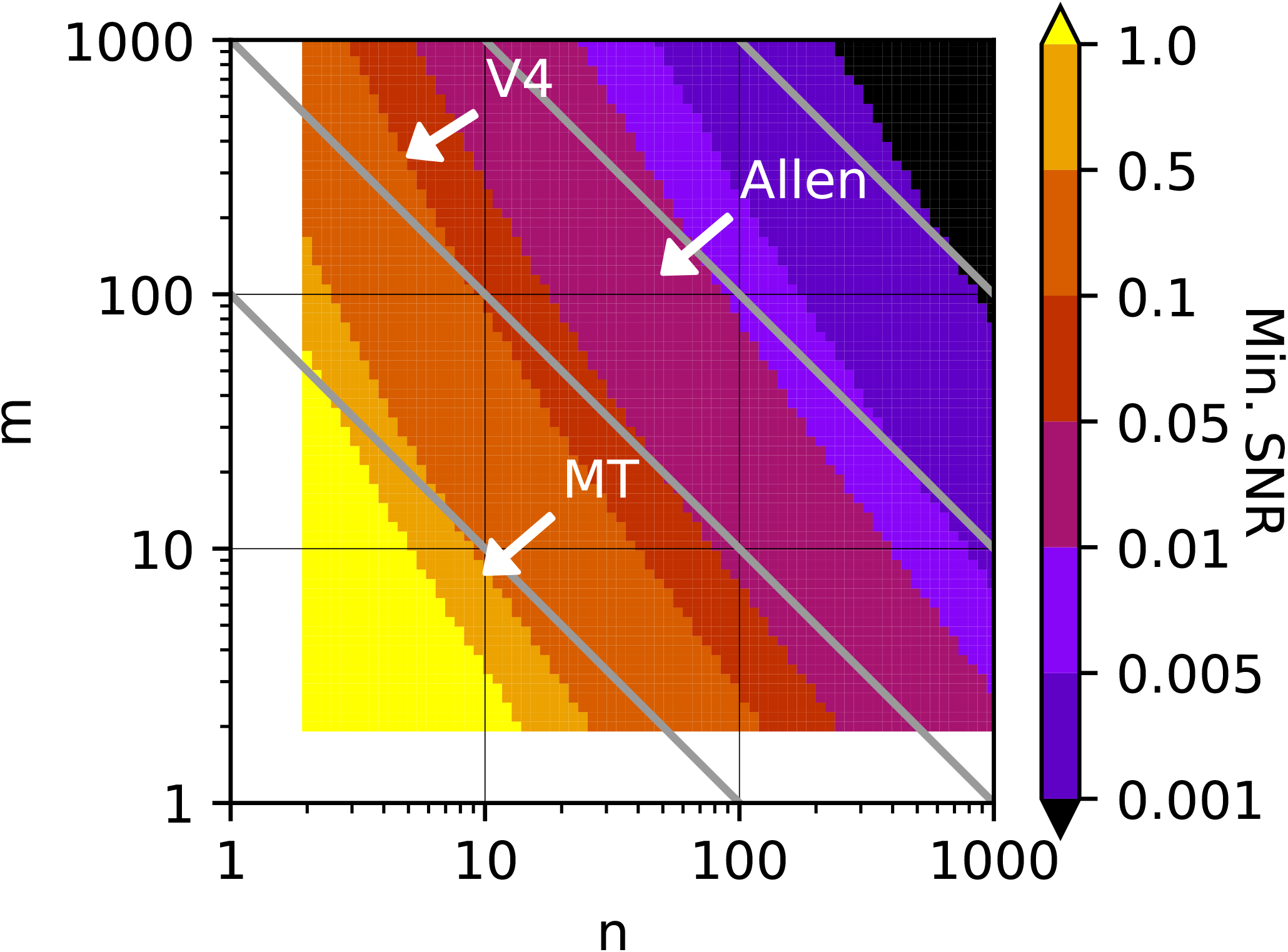
The minimal SNR needed to reliably detect tuning as a function of *m* the number of stimuli and *n* the number of trials. Approximate *m* and *n* are labeled for the datasets used in Figure 13.

On this map, we can locate points corresponding to the *m* and *n*, roughly, for data sets in Figure 13. The three V4 data sets have about the same number of stimuli and repeats (arrow marked “V4”, Figure 14), and thus require SNR≈0.1 or greater, implying that from 3% to 23% of the V4 data does not pass the criterion (Figure 13A, red and green traces, respectively, define endpoints). The MT data has the fewest number of total trials and thus has the highest threshold SNR≈0.5, which leaves 10% of the neurons with poorly estimated tuning curves. If more stimuli had been used at the expense of fewer repeats, say *n* = 2 and *m* = 40, then only a quarter of the neurons would have exceeded the increased threshold of SNR>1. The Allen Ca^2+^ and spike data sets both had similar *m* and *n*. Relative to the other data sets they had far more total trials and a greater number of repeats, thus the SNR criterion is substantially lower (SNR>0.01). Still, for the Ca^2+^ data, ~ 37% of the grating and ~ 25% of the natural image data did not have reliable tuning (Figure 13A, pink and grey thin trace). The Allen spiking data on the other hand had much higher SNR, and thus more trials could have been spent on expanding the stimulus set and fewer on repeated presentation (Figure 13A, thin brown purple trace).

## Discussion

### Summary

We have investigated the estimation of the correlation between a model prediction and the expected response (ER) of a neuron. It is established that trial-to-trial variability will cause the classic estimator, Pearson’s *r*^2^, to underestimate correlation, but there has been no direct comparison of prior methods to account for this confound. Our comparison of these methods revealed that some grossly over estimate correlation in high noise conditions. We built upon the best performing method to derive a more generally applicable estimator that we call 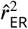 that performs as well or better than prior methods. We analytically validated this method by determining that it was a consistent estimator in the number of stimuli. We found in simulation it had a small upward bias, but that this was only appreciable at very high noise levels. None of the prior methods that we examined had validated confidence intervals, thus we developed confidence intervals for 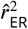. Motivated by the failure of generic bootstrap methods to achieve satisfactory confidence intervals, we developed a confidence interval method that outperformed them.

Applying our estimator to neural data, we demonstrated its essential value. In the case of MT recordings, it was able to unambiguously identify neurons for which a sinusoidal model was a good fit versus those for which it was a poor fit specifically because of systematic deviation and not because of noise. The associated confidence intervals allowed the systematic identification of noisy recordings that served no practical use in assessing the fit of the model. Poor model fits caused by noise vs. those caused by systematic differences in selectivity have very different interpretations, yet the traditional 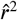 does not differentiate them while 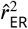 does.

Application of the estimator to the winning UW neural data challenge model, a DNN, provides the only validated assessment of state-of-the-art predictive model performance in V4. The estimator along with its CIs identified neurons that were challenging to the DNN and perhaps require a different modeling approach. It also validated the existence of single units that had nearly 50% of their variance explained, indicating that the DNN functionally captured a substantial part of what these units encode across natural images and thus could provide real insight into naturalistic V4 single unit encoding. On a practical level, we showed how the estimator allows for gains in trial efficiency since it converges more rapidly than 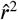 (Figure 3B and Figure 12). This is important when many stimuli are needed to validate models of high dimensional neural tuning.

Our tests on experimental data revealed that some neurons had confidence intervals covering the entire range of possible values, motivating us to propose the signal-to-noise ratio (SNR) as a metric of neural recording quality in the context of model fitting. We provide an unbiased estimator of SNR (15) and a practical interpretation: for a given number of stimuli and repeats, the SNR should be sufficient to reliably detect stimulus-driven response modulation on the basis of an F-test (Eqn. 21). We examined a variety of data sets and found (1) they differed with respect to how stimuli vs. repeats (m vs. n) were balanced, but adjustments can be made on the basis of SNR for guiding more efficient experiments (2) We found that there were large differences in SNR across data sets that are likely related to recording modality (e.g., Ca2+ imaging vs. electrode recording), which could be important for selecting appropriate experimental approaches and for guiding the refinement of current techniques to improve SNR.

### Interpretation of 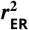

We have introduced an estimator and confidence intervals for the correlation between the true tuning curve of a neuron (its expected value across stimuli) and model predictions: 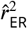. In the context of sensory neurophysiology, we believe it is reasonable to think of 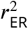 as reflecting solely how well a model explains a sensory representation. We justify this by the fact that 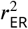 is solely a function of *E*[Response|Stimulus] thus solely a function of stimuli. We note two caveats: (1) non-sensory signals can influence sensory responses, e.g., eye movements which are non-random w.r.t stimuli and (2) *E*[Response|Stimulus] is not the only component of the sensory response, e.g., variability can be a function of stimuli (Cohen and Kohn, 2011).

### Relationship to prior methods

Here we followed a line of research (Sahani and Linden, 2003; Haefner and Cumming, 2009) that explicitly attempted to construct an unbiased estimator of *r*^2^ in the absence of noise or equivalently as *n* → ∞. Sahani and Linden constructed an unbiased estimator of the variation in the expected response of the neuron (i.e. tuning curve) they called this ‘signal power’. They normalize an estimator of explained variation, unbiased with respect to noise under conditions they did not specify (1-parameter regression of model predictions, see Appendix: Normalized Signal Power Explained (SPE)), and call this ‘normalized signal power explained’ (SPE). Sahani and Linden, by not specifying how a given model prediction should be fit to the neural data before estimating the quality of the fit, introduced potential problems in their estimator. This was recognized by Schoppe et al. (2016), who point out that the estimator was sensitive to differences between the mean of the model predictions and the neural data. Consequently, the estimator could give large negative values because the mean squared error between model predictions and neural responses was unbounded. This criticism while technically correct is easily overcome by regressing (with intercept term) the given model predictions onto the neural data before using normalized SPE. Schoppe et al., motivated by the problems they found in SPE, focused on simplifying CC_norm_ of Hsu et al. (2004) to not require re-sampling by making use of the signal power estimator developed by Sahani and Linden. They derived a simple estimator whose square we And is essentially numerically equivalent to the normalized SPE of Sahani and Linden in the case of 1 parameter regression. Since often the scaling and mean of a models predictions are irrelevant to questions of tuning, or deviation of scaling and mean of the model from data is better analyzed independently, it can be useful to understand an estimator of tuning similarity with respect to its performance when these are factored out.

Haefner and Cumming’s (2008) 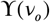 takes inspiration from Sahani and Linden but explicitly assumes that the model being fit is linear in its coefficients and so appropriately take into account degrees of freedom. We take a similar approach to Haefner and Cumming though provide one improvement: their formula requires the calculation of the sample variance because their derivation relies on the F-distribution formed by taking the ratio of the sum of squares of model residual over the sample variance (see Appendix: 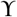). This in turn requires that the data have multiple repeats. There are two scenarios where this would be problematic. First, if stimuli are never repeated in an experiment (for example, in free viewing experiments), then one has to a priori assume trial-to-trial variability either from previous experiments or by assuming a known mean-variance relationship (e.g. the square root of Poisson distributed spiking gives 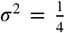). Second, if variance is stabilized by z-scoring each set of responses from the neuron, then the estimate of trial-to-trial variability is no longer a *χ*^2^ random variable but a constant *s*^2^ = 1. In our case, the estimate of trial-to-trial variability need not be *s*^2^, as long as it can be assumed to be unbiased. For example, it could be a constant if the experimentalist is willing to assume trial-to-trial variability is known.

Haefner and Cummings estimator differs from ours in two other primary ways: (1) they estimate fit quality through an unbiased estimator of the residual variance (unexplained variance), we instead estimate the variance of the predictor (explained variance), (2) their 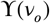 estimates variance explained for a linear model, ours estimates Pearson’s *r*^2^. Ours can be seen as a special case of theirs, being identical when estimating the fit of a linear model with a slope and intercept. We also provide the more general estimator of variance explained for a linear model (Eqn. 22) with the advantages discussed in the previous paragraph, but throughout this paper we deal with the simpler estimator.

### SNR

We found differences in SNR can be substantial and widely varying across neurons, data sets, and recording methods. Given the rise in large scale recordings and sharing of neuronal data, we believe unbiased estimates of SNR should be reported so that researchers can quickly judge whether a data set has sufficient statistical power and to aid in the design of comparable experiments. We provided concrete criteria by which to interpret SNR: the statistical power to detect stimulus-driven response modulation. Strikingly, in even a small sample of data sets, many neurons do not pass this criteria, suggesting that the adoption of a standard criterion for data quality, such as our SNR metric, could have a major impact in practice. Furthermore, guided by the metric the experimentalist can take steps to improve SNR by increasing stimulus duration and associated spike counting windows or customizing stimuli to the preferences of a neuron. On the other hand they can ameliorate the deleterious effects of low SNR by favoring repeats over number of stimuli (Figure 14).

One interpretation of the SNR metric we introduced is that it quantifies, for the time scale of the spike count window, the overall variance in the responses of the neuron attributable to the tuning curve of the neuron vs. trial-to-trial variability about that tuning curve. For example on the time scale of 1 second, many of the spiking recordings we examined had a large fraction of neurons with SNR > 1 meaning more variance was caused by the stimulus than by other sources (Figure 13B blue, orange, green traces median SNR>1). On the other hand, an appreciable number of neurons are dominated by their trial-to-trial variability, thus the tuning curve has less explanatory power about the responses of these neurons. Whether this is the result of stimulus choice and perhaps would be different for a more naturalistic setting is an open question. Recent theoretical and experimental work has argued that weakly tuned and untuned neurons can contribute to sensory encoding (Leavitt et al., 2017; Insanally et al., 2018; Zylberberg, 2018). The corrected estimate of SNR we provide (Eqn. 15) along with naturalistic stimulation can help to identify such neurons.

### Further work

Small improvements can be made to this estimator by decreasing its bias in the case of very low SNR (see Methods: Bias of 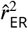). In the case of very low SNR, a single neuronal recording has little inferential power, but across a population of neurons, estimates of the average correlation to a model’s predictions can have low enough variance to provide useful inference. Yet, at very low SNR an appreciable bias begins to appear that will remain in the population average. One step to decrease this bias is to estimate the remaining bias (see Eqn 16) and use these estimates to perform a final adjustment of the estimate. We have not found a method which corrects this remaining bias at low SNR while keeping variance reasonable.

Here we have derived an estimator for the case where deterministic model predictions are correlated to a noisy signal. Often, one noisy signal is correlated to another for example when judging whether two neurons have similar tuning (termed signal correlation). We will show how the methods described in this paper can be extended to the neuron-to-neuron case in an upcoming publication (Pospisil and Bair, in prep).

A subtle but important point about our estimator is that it assumes stimuli are fixed: it estimates the 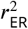 for the exact stimuli shown. An investigator may be interested in the quality of a model more generally, e.g., for a large corpus of natural images of which only a small fraction can be included in a recording session. In this case, to make population inferences the experimentalist can draw a random sample of images, collect neural responses, fit a model, then test that model on responses to a test set of images. In this case employing the random test set will account for over-fitting and using 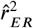 will account for noise in the neural responses in the fit to the validation set. Crucially though, the variability in the parameters of the model induced by the random training sample will not be accounted for. Intuitively, neural responses to one subset of images will give different estimated model parameters than another even in the absence of trial-to-trial variability. The correct interpretation of 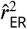 in this case is that it estimates how well a model can perform given finite noisy training data on noiseless validation data. It would be incorrect to interpret it as the best the model could possibly perform given infinite training data. Indeed, with more neural responses and less noise, model validation performance would improve. David and Gallant (2005) explore this issue calling it ‘estimation noise’ and provide an extrapolation method for estimating the fit of a linear model given unlimited stimuli. The estimator was not evaluated in terms of its bias or variance, and no analytic solutions that directly remove the bias of finite training data have been proposed. Both are valuable directions to pursue: the former to build confidence in the current method and the latter for potential gains in trial efficiency. A data driven re-sampling approach may be unavoidable when evaluating more complex models where the relationship between the amount of training samples and model performance would be analytically intractable, such as a DNN or biophysical model.

## Materials and Methods

### Simulation procedure

To simulate model-to-neuron fits, the square root of neural responses, *r_i,j_*, for the *i*th of *m* stimuli and the *j*th of n trials are modeled as independent normally distributed responses:

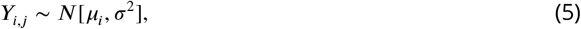

where variance *σ*^2^ is the same across all *Y_i,j_*. The mean response of the neuron to the *i*th stimulus (tuning curve) is 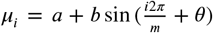 (Figure 1A, green trace solid dots) whose correlation to the model predictions 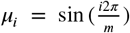 (red trace solid dots) are estimated, and the true correlation is 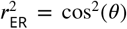. The results of the simulation are only a function of the magnitude of the vector of mean responses 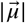 and the angle between the model predictions and true responses 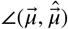. We choose a sinusoid for the simplicity of adjusting the phase, *θ*, to simulate different 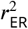.

From this model we draw *n* responses for each of the *m* stimuli (green open dots) and apply our estimator to this sample. We repeat this across many IID simulations to accumulate reliable statistics.

### Assumptions and terminology for derivation of unbiased estimators

Below we derive an unbiased estimator of the fraction of variance explained when a known signal is being fit to noisy neural responses. For this derivation, we assume the responses have undergone a variance stabilizing transform such that trial-to-trial variability is the same across all stimuli. For example, if the neural responses are Poisson distributed, *Y_i_* ~ *P*(*λ_i_*), where *Y_i_* is the response to the *i*th stimulus, which has expected response *λ_i_*, then a variance stabilizing transform is the square root. In particular, if 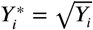, then,

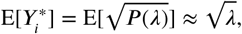

and

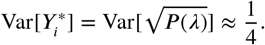

The mean of the transformed response, 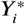, still increases with *λ*, whereas the variance is now approximately constant. To improve the estimate of the mean response, *n* repeats of each stimulus are collected. Invoking the central limit theorem, we can make the approximation:

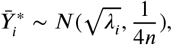

where the average across the *n* repeats is approximately normally distributed with variance decreasing with *n*. The assumption of a Poisson distributed neural responses is not always accurate. A more general mean-to-variance relationship,

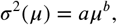

can be approximately stabilized to 1 by,

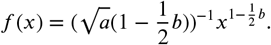

A square root will stabilize any linear mean-to-variance relationship, but unknown relationships require thattheir parameters be estimated. In the case of the linear relationship, this simply requires taking a square root and then, since the variance is constant across stimuli, averaging the estimated variance across all stimuli. If it is not reasonable to assume a parametric mean-to-variance relationship and there are enough repeats, one can simply divide all responses to a given stimulus by their sample standard deviation to achieve *σ*^2^ ≈ 1. For the derivation below, we assume that variance-stabilized responses to *n* repeats have been averaged for each of *m* stimuli to yield the mean response to the i^th^ stimulus: 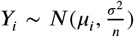, where *σ*^2^ is the trial-to-trial variability and *μ_i_* the *i*th expected value.

### Unbiased estimation of *r*^2^

Given a set of mean neural responses *Y_i_* and model predictions *v_i_* the naive estimator 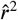 is calculated as follows:

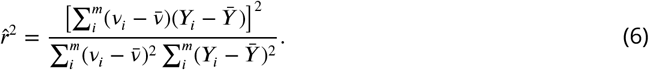

Our goal is to find an estimator such that,

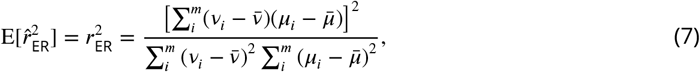

where 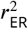 is the correlation in the absence of noise, i.e., the fraction of variance explained by the model prediction, *v*, of the expected response (ER), *μ_i_*, of the neuron. Our strategy will be to remove the bias in the numerator and denominator separately and then reform the ratio of these unbiased estimators for an approximately unbiased estimator.

### Unbiased estimate of numerator

The numerator of Eqn. 6, which we call 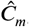, is a weighted sum of normal random variables that is then squared, thus it has a scaled non-central chi-squared distribution:

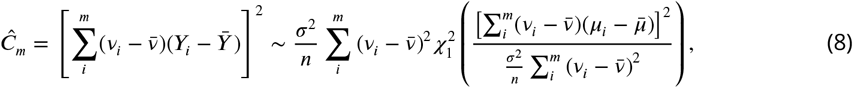

and its expectation is:

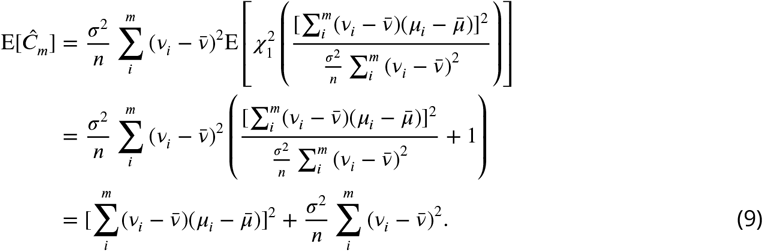

The term on the left in the final line is the desired numerator and the term on the right the bias contributed by *σ*^2^. To form our estimator, 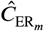, for the numerator of Eqn. 6, we simply subtract an unbiased estimator of this bias term:

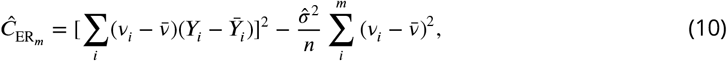

where 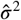 is typically the sample variance, *s*^2^, estimated from the data, but it can be any unbiased estimator even an assumed constant. For example, if stimuli are not repeated (i.e., *n* = 1) and one is willing to assume that responses are Poisson distributed, then the square root of these responses will give 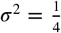 and thus one can substitute 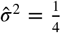. The case for the denominator is similar.

### Unbiased estimate of denominator

The denominator of Eqn. 6, which we call 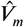, is a weighted sum of squared normal random variables and thus also follows a scaled non-central chi-squared distribution:

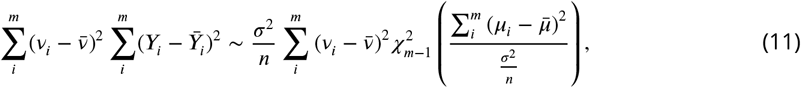

with expectation,

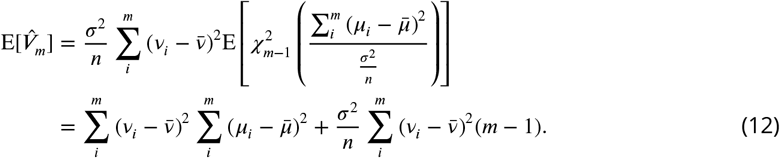

Similarly to the numerator, we find that the first term on the final line is the desired denominator, and the second term is the bias. Thus we subtract off an unbiased estimator of the bias:

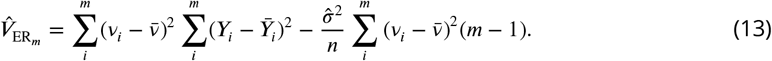

Taking the ratio of these two unbiased estimators we have:

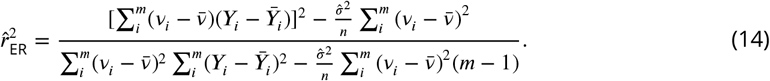

This equation can be further simplified by scaling the model predictions such that 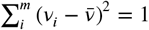.

### Estimators of correction terms

Two important parameters, 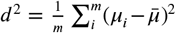 and *σ*^2^, are unknown. Below we provide unbiased estimators of each of these terms. An unbiased estimate of sample variance for trials of the *i*th stimulus is 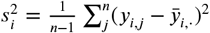 and then assuming the variance is the same across stimuli we can average over *i* for a global estimate:

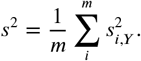

Throughout the paper we use this as our estimate of trial-to-trial variability 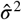.

For *d*^2^ we have:

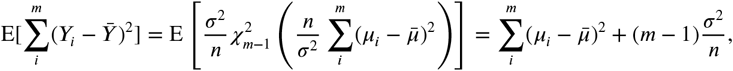

so an unbiased estimator is

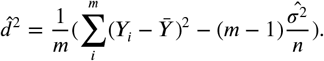

We use this estimator to correct the estimate of SNR (4) for trial-to-trial variability as follows:

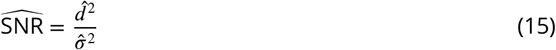

### Bias of 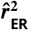

To remove the bias of Pearson’s 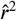 we follow the approach of subtracting off its effect in the numerator and denominator. Prior work has not examined the potential problem with this approach: the expectation of a non-linear transformation of a set of random variables is not necessarily the transformation of their expected values. In this particular case the expectation of the ratio is not necessarily the ratio of the expectations: 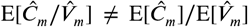 (where we have let 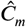 be the unbiased numerator and 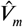 the unbiased denominator). Thus even though we have removed the bias in the numerator and denominator it does not imply their ratio is unbiased. Calculating the expectation of the ratio we see the conditions under which it will be unbiased:

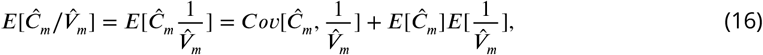

so 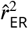 is unbiased if 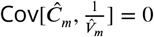 and 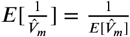 but we find in simulation often 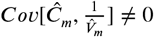 and by Jensen’s inequality 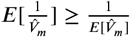.

Thus if the estimator 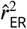 is not unbiased for 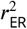 what recommends it over the naive 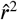? While we mainly focused on how in simulation for typical ranges of parameters it has a lower bias (Figure 3) it also has a theoretical justification. As we saw in simulation as *m* the number of stimuli increases, it becomes closer to being unbiased while 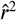 does not (Figure 3C). Convergence to the parameter of interest, otherwise known as consistency, gives a theoretical justification for an estimator. Below we show that 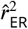 is consistent for 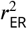 while 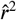 is not.

We note that further refinements of 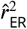 could involve estimating 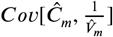 and the difference between 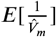 and 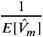 via simulation then correcting for them (see Discussion).

### Consistency of 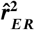 in *m*

We aim to show that 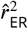 is consistent for 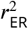 in *m*, more formally:

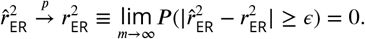

We make use of the continuous mapping theorem that guarantees if a random vector 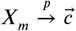 then for a continuous transformation 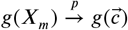. Taking our random vector to be our estimates of the numerator (Eqn. 10 and denominator (Eqn. 13), 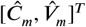, and our continuous transformation to be forming the ratio, 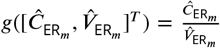, all we need to show is that 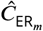 and 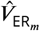 themselves are consistent estimators for the numerator and denominator of 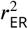 then their ratio will be consistent for 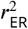.

First, we have already shown that 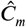 and 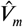 are unbiased estimators. Next we must show that their variance is decreasing with *m* then via Chebyshev’s inequality:

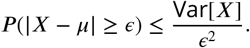

Here we consider the case where 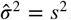. Since the model predictions (*v_i_*) are fixed for the purpose of the proof, we assume the dot product between model predictions and neural responses is scaled linearly by *m*

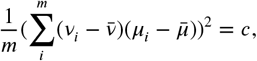

as is the dynamic range of the neuron:

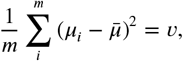

and we scale the numerator and denominator by 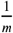 which makes no change to their ratio.

The numerator: 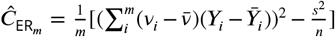 has variance equal to the sum of the variance of its first and second term (since they are independent). The variances are, respectively,

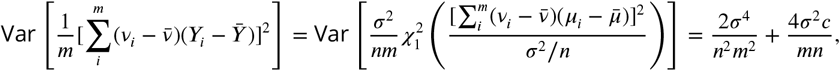

and

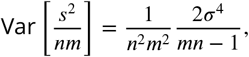

thus

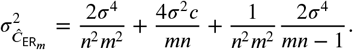

The denominator, 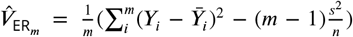, also has variance equal to the sum of the variance of its first and second term (by independence). The variances are, respectively,

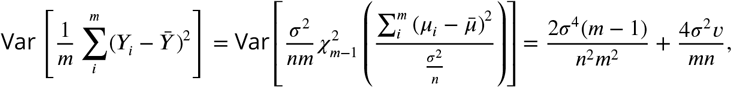

and

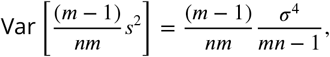

thus

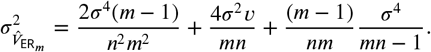

Noticing that for both 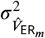 and 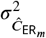 all but *m* is constant we can find an *m* to scale variance below any given *ϵ*. So by Chebyshev’s inequality we have

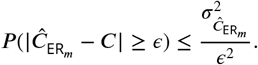

Since as 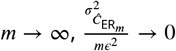 we have that:

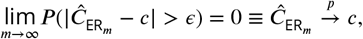

and similarly

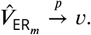

Thus by the continuous mapping theorem:

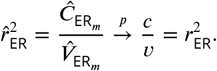

In contrast we show below that the naive estimator is not consistent and provide insight into when the difference between 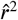 and 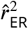 is large.

### Inconsistency of 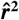 in *m*

Similarly to the previous derivation, we can take the numerator and denominator of 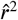 (Eqn. 8, 11), scale by 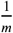, then find their asymptotic values, to in turn find the asymptotic value of 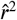 though we simplify by setting the model to be unit length:

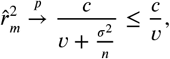

thus 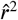 is not a consistent estimator (in *m*) of 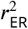.

When will the bias of 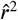 be large? Here we consider this when 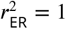. In this case:

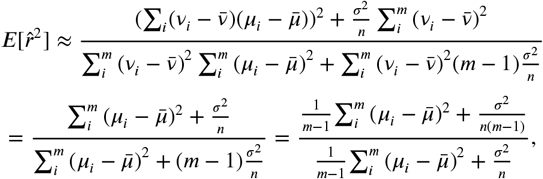

where we have simply formed the ratio of the expectation of the numerator and denominator (Eqn. 9, 12) and set the length of the model predictions to 1. We then switch to average dynamic range in the second equality (note 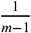 divided into numerator and denominator). This makes it clear that while the numerator is getting closer to its true value with *m* the denominator is fixed to the ratio determined by its average dynamic range and trial-to-trial variability. Thus *r*^2^ is always stuck between the two ratios:

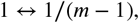

going towards the term on the left as the dynamic range increases and the term on the right as trial-to-trial variability increases. Thus increasing *m* will not decrease the bias (it will increase it but only slightly for large *m*) only increasing *n* and dynamic range or decreasing trial-to-trial variability will decrease the bias and *n* and *m* are the only parameters truly under experimenter control.

### Quantifying uncertainty in the estimator

#### Confidence Intervals for 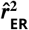

Here we develop a method that we prove provides *α* level confidence intervals forthe estimator 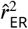. We considered typical approaches (the parametric bootstrap and non-parametric bootstrap), but found that they were not reliable for typical ranges of parameters (see Results: Confidence intervals for 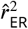.

Our approach in essence is we find the lowest 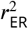 whose distribution would give an estimate greater than the observed 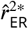 with probability *α*/2 calling this 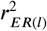 and the highest 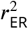 that would give an estimate less then the observed 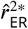 with probability *α*/2 calling this 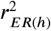. The interval 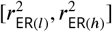 then serves as our *α*-level confidence interval (see Figure 15 for graphical explanation). We use a Bayesian framework to sample from the distribution of 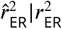 given the observed neural statistics *s*^2^ and 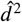 that allows us to find 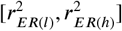 (see Computing confidence intervals, below) under assumed uninformative priors of the parameters *σ*^2^ and *d*^2^.

**Figure 15.**
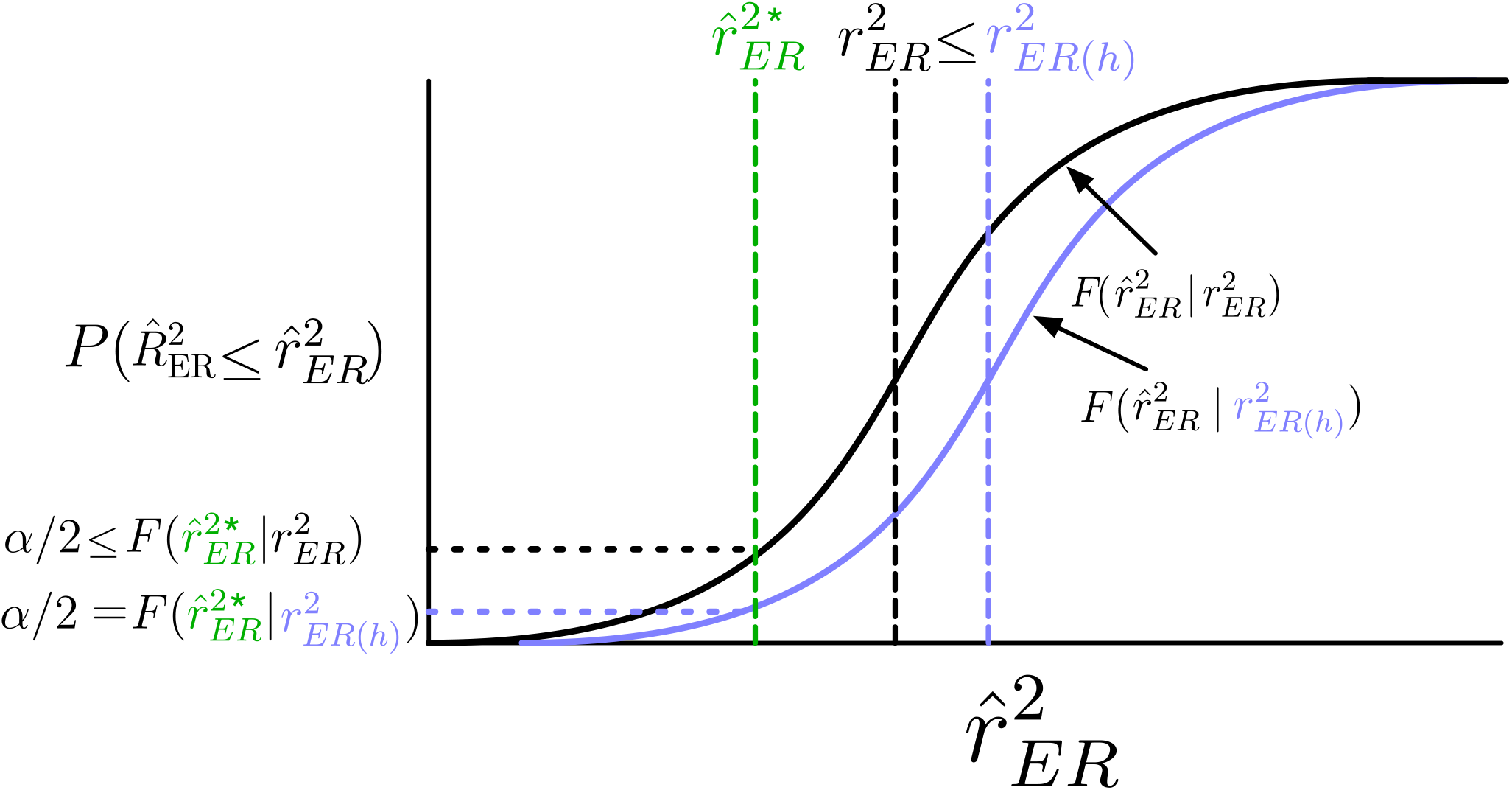
Illustrative schematic of confidence interval estimation. Given a sample (green dashed vertical) from the distribution 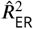 (cumulative distribution solid black curve) associated with the true 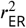 (black dashed vertical) the upper limit of the *α* level confidence interval is the 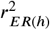 (purple vertical dashed) that would generate 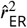 less than the observed with probability *α*/2 (purple horizontal dashed). Under the assumption 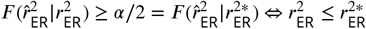 the event that 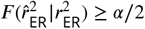 corresponds to the event that 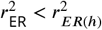 thus the upper limit of the confidence interval contains the true value of 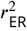. In graphical terms if the black horizontal dashed line is above the purple then it is guaranteed the vertical dashed purple is in front of the black horizontal dashed. Thus these two events have the same probability: 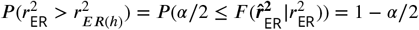.

Here, we justify this procedure for the case of 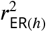 (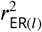 is similar). Our two main assumption are that the distribution 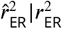 is stochastically increasing in 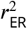:

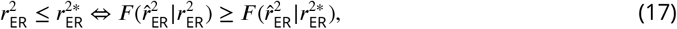

and that we can always find an 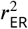 where for any observed 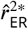:

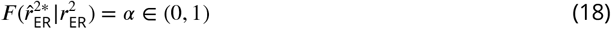

where *F* is the cumulative probability distribution. We then consider the two mutually exclusive possibilities: First, with probability 1 - *α*/2, the observed 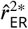 is large enough to satisfy:

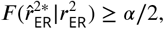

then by the assumption in Eqn. 18 we can find a 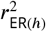 where:

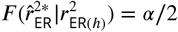

under our initial assumption (Eqn. 17) this implies

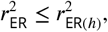

because

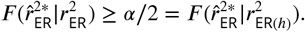

Second, if on the other hand 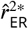 is small enough, with probability *α*/2, such that:

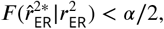

then:

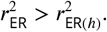

Thus, under repeated sampling the upper limit of our confidence interval, 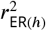, does not contain 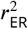 with probability *α*/2 as desired. The proof for the lower end of the confidence interval 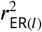 is similar. The probability of the mutually exclusive events that either 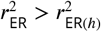 or 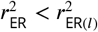 is the sum of the probability of the two events, *α*. See Figure 15 for a graphical explanation of this proof.

For simplicity of the proof we assumed we would always be able to find 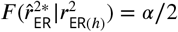 which is not necessarily the case because 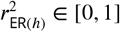 is bounded but 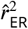 is not. If 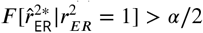 or 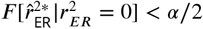 then there is no 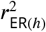 that will achieve *α*/2.

Under the condition where 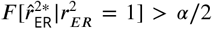 we simply set 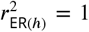 and since 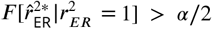 implies 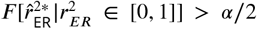 the confidence interval should contain the true value which is the case since 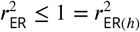.

Under the condition where 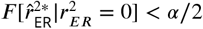 we set 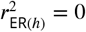 but we must place a seemingly strange requirement in this case and only because of the possibility that 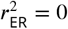. Here we must set the confidence interval, though normally inclusive to be non-inclusive. Intuitively this is because if 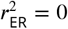 the upper end of the confidence interval would always contain the true value. In the other case, 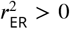, then similarly to above since 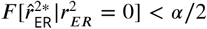 implies 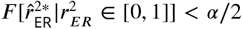 the confidence interval should not contain the true value which is the case since 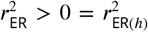. The case for 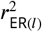 is similar.

In summary our confidence interval is defined to be 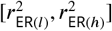 when 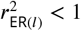 and 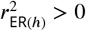 but 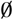 the empty set if 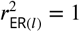 or 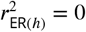. The lower interval 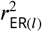 satisfies 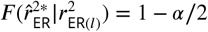 except if 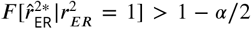 or 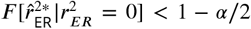 then respectively 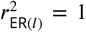 or 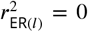. The upper interval 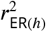 satisfies 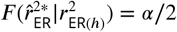 except if 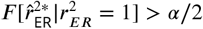 or 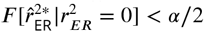 then respectively 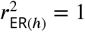 or 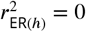.

To sample from the distribution of 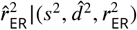 we assume that *σ*^2^ and *d*^2^ follow an uninformative non-negative uniform prior and given the observed *s*^2^ and 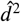 we obtain samples from the posterior distribution of *σ*^2^ and *d*^2^ via MCMC sampling (for details see Bayesian Sampling). Finally for a chosen 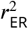 (e.g. a candidate for 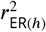) we sample from 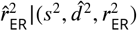 by drawing *σ*^2^, *d*^2^ samples from the posterior 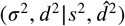 while 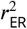 is fixed to the desired value thus for each sample we then draw observations *Y* and model *x* from the model described in Eqn. 5 and finally calculate 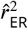.

#### Computing confidence intervals

We have described how we can sample from 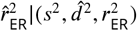 the distribution of our estimator under the assumptions described above. Here we describe how using these samples we find our confidence intervals 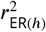 and 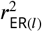. For simplicity we will describe the algorithm for 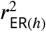. The algorithm uses a simple iterative bracketing to narrow down the range of candidate values for 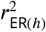 from [0,1] down to some arbitrarily small interval. First it evaluates the highest and lowest possible values for 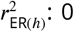 and 1. For example to evaluate 1 it does so by drawing samples from 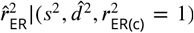(we draw 2,500) then calculating the number which were equal to or less then the original observed 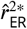 we find the estimated proportion 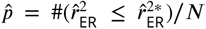 where *N* is the number samples drawn. We then calculate a z-statistic to test whether his number is equal to the desired *α*/2:

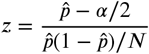

then at some desired significance level (here we use *p* < 0.01) we either do not reject the null and accept 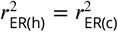 or if we reject and *z* is positive we determine that 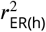 must be higher and if *z* is negative it must be lower. In the case where 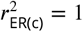 and *z* is positive there are higher possible values of 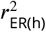 and thus we accept 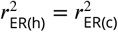. If instead we continue on to the next step we choose a new candidate by sampling from 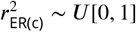 then evaluating the result and if we reject the null and *z* is positive our new interval will be 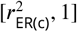 and if *z* is negative 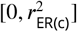 otherwise if we do not reject the null we accept 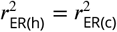. We continue this bracketing until we do not reject the null or a pre-determined number of splits has passed (here we use 100). Accuracy of this algorithm will increase with number of splits and the number of simulation samples.

#### Confidence interval validation

We evaluate the confidence intervals under the sampling distribution 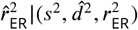. Conceptually this is the distribution 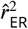 where data has already been collected and sample variance and sample dynamic range calculated and now we wish to calculate the data’s fit to a model with unknown but fixed 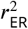. To demonstrate our method in simulation, for proof see above, and that it contains the unknown 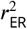 at the desire proportion *α* we follow this procedure: for a chosen n, m, *σ*^2^, *d*^2^, and 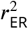 we first sample an *n* ×*m* data matrix (Y) and for this data we calculate the sample variance *s*^2^ and dynamic range 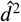. Then using the Metropolis-Hastings algorithm described we draw 5,000 samples from the posterior 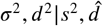. We now simulate the distribution of 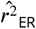 from this data from the given 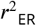 drawing 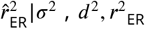 where|*σ*^2^, *d*^2^ are being drawn from the aformentioned posterior and 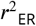 is given and thus fixed. For each of these draws we construct confidence intervals. Finally we calculate the proportion of times which the confidence intervals contained the true 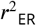 as an estimate of the true *α* level of the confidence interval method.

#### Bayesian model and simulation

We sample from the posterior of two parameters: 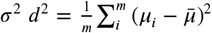. Their associated sufficient statistics are:

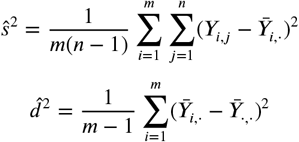

and their distributions are:

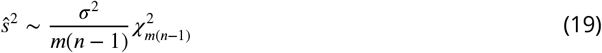

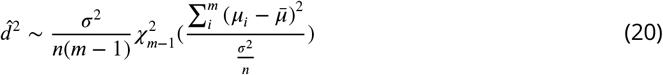

By Bayes theorem we have that:

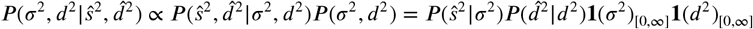

where the final equality is derived by recognizing the sample variance 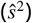 and dynamic range 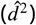 are independent and setting the prior to be uniform non-negative. The estimates 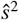 and 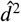 are fixed, calculated from the data, and our goal is to look up the distribution of the parameters given these fixed values. We use the Metropolis-Hastings algorithm to draw from the desired distribution 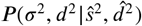 and approximate it with the empirical distribution (a histogram). Our sampling procedure is as follows: we initialize our parameter samples *σ*^2^, *d*^2^ at their estimates 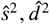 then from our proposal distribution a truncated multivariate normal with means 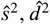 and diagonal variances equal to the variance of the distributions (Eqns. 19 and 20) where 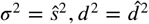 we sample a new candidate. We take the ratio of likelihoods 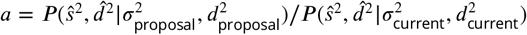. If the ratio is greater than 1 we accept the candidates as our new current samples and if it is less than 1 we then draw from *u* ~ *U* [0,1] and if *u* < *α* we also accept the candidates but if not we retain the current samples. Throughout the paper we run the chain for 5,000 iterations, then when using the trace we randomly sample with replacement from them.

#### SNR relation to F-test and number of trials

Our goal is to be able find for a given SNR and number of repeats the number of stimuli needed to reliably detect tuning under an F-test. To calculate the F-statistic for testing whether there is variation in the expected responses across stimuli (i.e. some selectivity) we form the ratio:

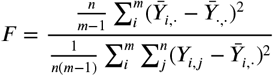

where the numerator calculates the amount of variance explained by stimuli and the denominator calculates the amount of variance unexplained by stimuli. The numerator is a scaled non-central *χ*^2^ distribution:

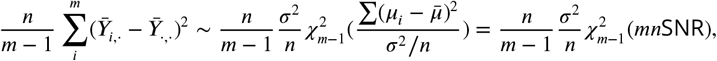

where the final equality comes from the definition of SNR (4). The denominator is a central *χ*^2^ distribution:

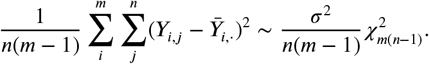

Thus taking the ratio we have a singly non-central F-distribution:

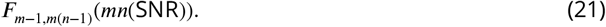

To test whether their is significant tuning we then set an alpha level criteria *c* under the null hypothesis that it is a central F-distribtion:

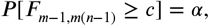

finally given that we know *m* and *n*, we can find SNR where for some high probability *β*

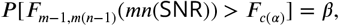

in this paper we set *β* = 0.99 and *α* = 0.01. We numerically solve for SNR in the equation above.

#### Electrophysiological data

We re-used a variety of neuronal data from previous studies. Experimental protocols for all studies are described in detail in the original publications.

We re-analyzed data from three previous single-unit extracellular studies of parafoveal V4 neurons in the awake fixating rhesus monkey (Macaca mulattta). Data from the first study, Pasupathy and Connor (2001), consists of the responses of 109 V4 neurons to a set of 362 shapes. There were typically 3-5 repeats of each stimulus and we used only the 96 cells that had 4 repeats for all stimuli. We used the spike count for each trial during the 500 ms stimulus presentation. To estimate translation invariance, we used data from a second study, El-Shamayleh and Pasupathy (2016). The data from the second study consists of responses of 39 neurons tested for translation invariance. The stimuli were the same types of shapes as the first study, but where the position of the stimuli within the RF was also varied. Each neuron was tested with up to 56 shapes (some of which are rotations of others) presented at 3-5 positions within the RF. Each unique combination of stimulus and RF position was presented for 5-16 repeats, and spike counts were averaged over the 300 ms stimulus presentation. To estimate fill-outline invariance we used data from the study of Popovkina et al. (2019). Filled stimuli were drawn from the same set as described for the previous two studies and outline stimuli were the same except the fill was set to be equivalent to background color and the outline width was set to 2,3, or 4 pixels (0.05-0.1 deg) with thicker outlines for more eccentric RFs. All stimuli color and luminance were customized to elicit a robust response from the recorded neuron. 43 well-isolated neurons were recorded from (7 from one monkey and 36 from the second). Spikes were counted over the 300-ms duration of each stimulus presentation.

MT data was recorded from three awake rhesus monkey (Macaca mulattta) viewing dynamic random dots. Stimuli is described in Britten et al., 1992. Optimal speed of drifting dots was found for the one of the two neurons being recorded. 8 different direction of motion at 45 deg increments were repeated 10-20 times. Monkeys performed a two alternative forced choice task of discrimination of motion direction task during the experiment. Post-stimulus spikes were counted in the 2 second window of stimulus presentation. Experimenters were rigorous in only recording from pairs of neurons whose spike wave forms were strikingly different.

Responses to natural images in V4 came from the V4 UW neural data challenge 2019. Up to 601 images were shown with between 3-20 repeats for each image. These images were drawn semi-randomly from the 2012 ILSVRC validation set of images where an 80X80 pixel patch was sampled then had soft window applied (circular alpha Gaussian with standard deviation of 16 pixels). There are two basic categories of recordings: multi-unit where the number of spikes from a small population of neurons is recorded and single unit where through careful post-processing it was insured the spikes recorded were from a single neuron. Multi-unit is essentially all of the detected spikes which did not come from the isolated single neuron. Images were shown for 300 ms with 250 ms in between images. The model we analyze was the winner of the Neural Data Challenge (out of 32 competitors) on held-out data from the 14 sets of V4 responses to natural images.

Detailed description of the mouse calcium data are given in Allen Brain Observatory (2016) and Devries et al. (2018) here we briefly summarize. Fluorescence of mouse visual cortex neurons expressing GCaMP6f was measured via 2-photon imaging through a cranial window. In the analyses here we examined signals recorded in response to natural scenes and static gratings. These images were presented for 0.25 seconds each with no interval between with 50 repeats in random order. The natural scene stimulus consisted of 118 natural images from a variety of databases. The static grating stimulus consisted of a full field static sinusoidal grating at a single contrast (80%). The grating was presented at 6 different orientations, 5 spatial frequencies (0.02, 0.04, 0.08, 0.16, 0.32 cycles/degree), and 4 phases (0, 0.25, 0.5, 0.75).

For every trial the Δ*F*/*F* was estimated use the average fluorescence of the preceding second (4 image presentations) as baseline. The ‘response’ we used in our analysis was the average change in fluorescence during the 1/2 second period after the start of the image presentation relative to 1 second before.

The Allen mouse neuropixel SUA data was recorded in response to the same stimuli as the calcium data described above (Siegle et al., 2019). Spike counting windows were 0.25s.

## Appendix

### Prior analytic methods of estimating 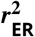

Here we write out the exact formulas of two prior methods (Sahani and Linden 2003; Haefner and Cumming, 2009) that are closely related to 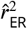 in the notation we have used through out the paper for the purpose of comparison.

Our method is derived via the strategy of Haefner and Cumming to unbias the numerator and denominator of the coefficient of determination. Haefner and Cumming in turn cite Sahani and Linden as the predecessor to their method. Both papers sought an estimator of the quality of model fit unbiased by noise. Haefner and Cumming explicitly develop an estimator of the fraction of variance explained also known as the coefficient of determination. If the model is linear (in its coefficients), fit by least squares, and an intercept is fit (or equivalently the mean of the data is subtracted off) the coefficient of determination is equivalent to using Pearson’s *r*^2^. We focus on the case when these conditions are true as the estimator is amenable to direct analysis in particular accounting for degrees of freedom is straightforward for linear models but not non-linear models. Below we first work out the asymptotically (in *m*) expected value of the estimator ‘Normalized Signal Power Explained’ (SPE) of Sahani and Linden and then the estimator 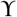 of Haefner and Cumming.

For all estimators below we assume responses are to *m* stimuli with *n* trials whose variance have been stabilized. The response to the *i*th stimuli and the *j*th trial is *Y_i,j_* ~ *N*(*μ_i_, σ*^2^) where *σ*^2^ is the trial-to-trial variability and *μ_i_* the *i*th expected value of responses post variance stabilizing transform. The predictions are fixed for the *m* stimuli and the *i*th predicted expected value of the data is *v_i_*. When averaging data across trials our notation will be 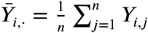 and across stimuli 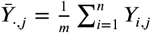.

### Normalized Signal Power Explained (SPE)

In the original publication describing SPE Sahani and Linden do not unambigously specify how to calculate it. In particular they do not specify when calculating sample estimates of what they call ‘power’, but is better known as variance, whether they normalize by *m*–1 (the unbiased estimate) or *m* (the MLE) or when they calculate the difference between variance of the measured response and the residual (section 3 first paragraph) they do not specify whether this is for the average across trials. We thus use the formula as described by Schoppe et al. who provide code to calculate SPE thus specify the formula unambiguously (though it differs from the formula they write in their paper in particular their 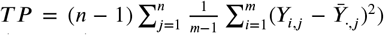 (Eqn. 5 of Sahani and Linden) but in their code it is 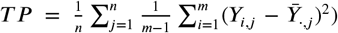, which in the context of the quantities being estimated, makes more sense (see calculation of expected value of denominator). For comparison to the derivation in Schoppe we give their notation and its equivalent terms in our notation: *N* = *n*, *T* = *m*, *R* = *Y_i,j_*, 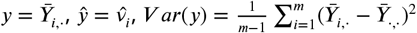, and finally their estimator:

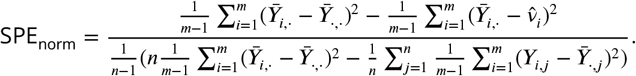

Calculating the expectation of the numerator and the denominator we can find the asymptotic expectation. Numerator:

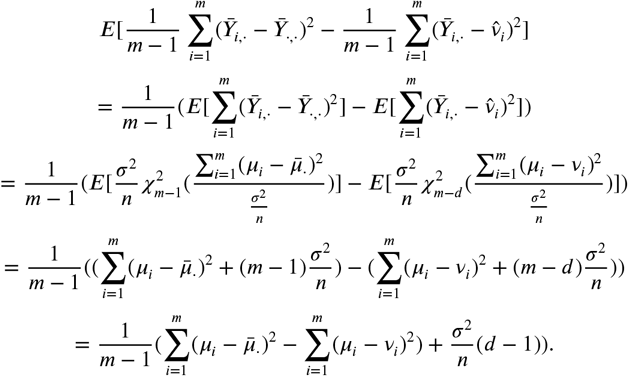

Denominator:

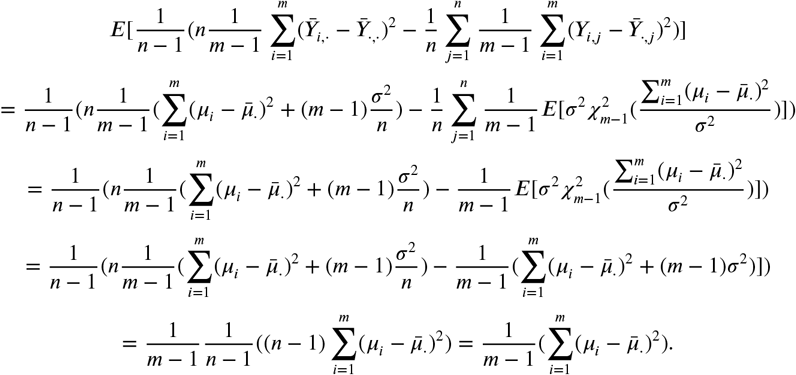

Putting the expectations into the numerator and denominator we have:

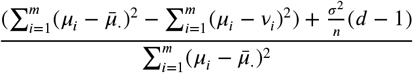

we note that only if *d* = 1 (i.e. the model has only 1 term) is the numerator unbiased which was the motivation for Haefner and Cumming to develop an estimator that accounts for degrees of freedom of a linear model.

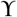

For comparison to their original paper we give their notation and its equivalent term in our notation: *d_i,j_* = *Y_i,j_*, 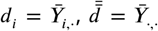, Σ^2^ = *σ*^2^, *N* = *m*, *N_σ_* = *m*(*n* – 1), *R* = *n*, *n* = *d*, *D_i_*, = *μ_i_*, *M_i_* = *v_i_*, 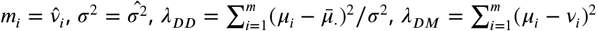.

These authors explicitly attempt to remove the bias of the coefficient of determination:

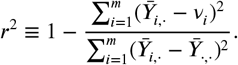

Their unbiased estimator is derived by dividing the numerator and denominator by the sample trial-to-trial variability 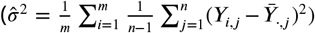 estimate and their respective degrees of freedom (*d* below being the degrees of freedom of the linear model) and noting the two terms become non-central F-distributions then shifting and scaling these random variables so they provide unbiased estimates of the numerator

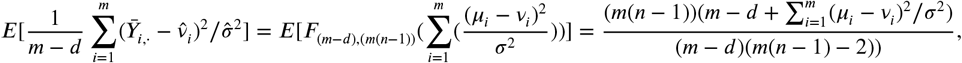

thus the unbiased estimate of the numerator is:

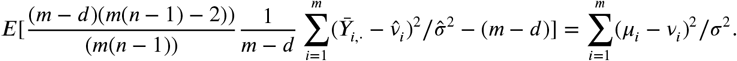

The expectation of the denominator is:

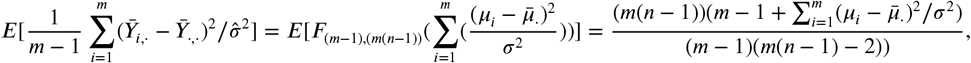

thus the unbiased estimate of the denominator is:

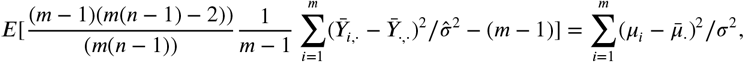

plugging these into the ratio we have their estimator:

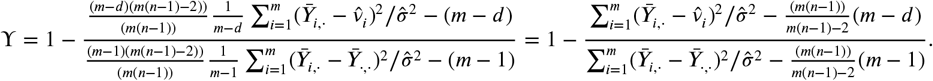

It should be noted they state that the final estimators numerator and denominator (Eqn. 8 of Haefner and Cumming) are unbiased but this is not true. The final algebraic simplification, the final term on the right, does not have a numerator and denominator unbiased for 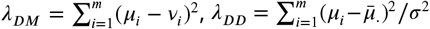, though the ratio is equivalentto the term on the right which does have an unbiased numerator and denominator. It should also be noted that the derivation via the non-central *F* was not necessary: the expectation of the numerator and denominator are straightforward to calculate as non-central *χ*^2^ random variables. While the term we derive 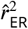 is explicitly meant to be the analogue to *r*^2^ where one plugs in two vectors of measurements we re-derive the Haefner and Cumming formula for the case when one wants to measure the strength of the linear relationship with the fit of a linear model. We diverge by not using non-central F-distributions so that it is not necessary to estimate variance to make the calculation (if for example there is a strong prior for the variance and/or multiple trials were not collected). Here we assume, as did Haefner and Cumming implicitly by using the non-central F, that *v_i_* were fit from a linear model via least squares with *d* coefficients. The expectation of the numerator is:

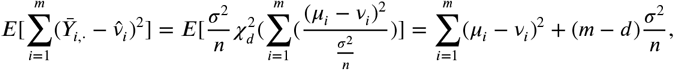

thus the unbiased estimate of the numerator is:

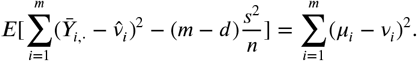

The expectation of the denominator is:

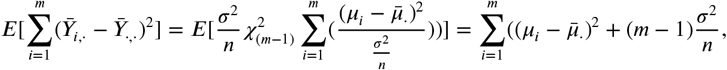

thus the unbiased estimate of the denominator is:

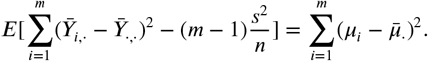

Thus their ratio forms:

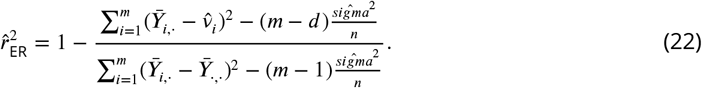

## Acknowledgments

We thank Greg Horwitz and Yen-Chi Chen for helpful suggestions and advice. We thank the participants in the UW neural data challenge including the winner Oleg Polosin for their neural models. The competition was funded by the UW Computational Neuroscience Center. This work was funded by a National Science Foundation (NSF) Graduate Research Fellowship (Pospisil), NIH/NEI R01-EY029997 and NIH/NEI R01-EY027023 (Bair).

## References

Allen Brain Observatory (2016) http://observatory.brain-map.org/visualcoding/. Accessed: 2019-506 04-20.

Bair WI, Zohary E, Newsome WT (2001) Correlated firing in macaque visual area MT: Time scales and relationship to behavior. TheJournal of Neuroscience: The Official Journal of the Society for Neuroscience, 21(5), 1676–1697.

Cohen MR, Kohn A (2011) Measuring and interpreting neuronal correlations. Nature Neuroscience 14:811–819.

Cadena SA, Denfield GH, Walker EY, Gatys LA, Tolias AS, Bethge M, Ecker AS (2019) Deep convolutional models improve predictions of macaque V1 responses to natural images. PLOS Computational Biology, 15(4), e1006897.

Siegle JH, Jia X, Durand S, Gale S, Bennett C, Graddis N, Heller G, Ramirez TK, Choi H, Luviano JA, Groblewski PA, Ahmed R, Arkhipov A, Bernard A, Billeh YN, Brown D, Buice MA, Cain N, Caldejon S, … Koch C (2019) A survey of spiking activity reveals a functional hierarchy of mouse corticothalamic visual areas. BioRxiv, 805010. https://doi.org/10.1101/805010

David SV, Gallant JL (2005) Predicting neuronal responses during natural vision. Network (Bristol, England), 16(2-3), 239–260.

de Vries SEJ, Lecoq J, Buice MA, Groblewski PA, Ocker GK, Oliver M, Feng D, Cain N, Ledochowitsch P, Millman D, et al. (2018) A large-scale, standardized physiological survey reveals higher order coding throughout the mouse visual cortex. BioRxiv.

Efron B, Tibshirani RJ (1994) An Introduction to the Bootstrap. CRC Press.

Haefner RM, Cumming BG (2009) An improved estimator of Variance Explained in the presence of noise. Advances in Neural Information Processing Systems 21 (pp. 585–592).

Insanally MN, Carcea I, Field RE, Rodgers CC, DePasquale B, Rajan K, DeWeese MR, Albanna BF, Froemke RC (2018) Nominally non-responsive frontal and sensory cortical cells encode task-relevant variables via ensemble consensus-building. BioRxiv, 347617.

Leavitt ML, Pieper F, Sachs AJ, Martinez-Trujillo JC (2017) Correlated variability modifies working memory fidelity in primate prefrontal neuronal ensembles. Proceedings of the National Academy of Sciences of the United States of America, 114(12), E2494–E2503.

Pasupathy A, Connor CE (2001) Shape Representation in Area V4: Position-Specific Tuning for Boundary Conformation. Journal of Neurophysiology, 86(5), 2505–2519.

Popovkina DV, Bair WI, Pasupathy A (2019) Modeling diverse responses to filled and outline shapes in macaque V4. Journal of Neurophysiology, 121(3), 1059–1077.

Pospisil DA, Bair WI (in prep) The unbiased estimation of *r*^2^ between neural responses.

Sahani MI, Linden JF (2003) How Linear are Auditory Cortical Responses? Advances in Neural Information Processing Systems 15 (pp. 125–132).

Schoppe O, Harper NS, Willmore BDB, King AJ, Schnupp JWH (2016) Measuring the Performance of Neural Models. Frontiers in Computational Neuroscience, 10.

Yamins DLK, Hong H, Cadieu CF, Solomon EA, Seibert D, DiCarlo JJ (2014) Performance-optimized hierarchical models predict neural responses in higher visual cortex. Proceedings of the National Academy of Sciences of the United States of America, 111(23), 8619–8624.

Zohary E, Shadlen MN, Newsome WT (1994) Correlated neuronal discharge rate and its implications for psychophysical performance. Nature 370:140–143.

Zylberberg J (2018) The role of untuned neurons in sensory information coding. BioRxiv, 134379.

